# Characterization of Histone Lysine β-hydroxybutyrylation in bovine tissues, cells, and cumulus-oocyte complexes

**DOI:** 10.1101/2021.09.07.459289

**Authors:** Juliano Rodrigues Sangalli, Ricardo Perecin Nociti, Maite del Collado, Rafael Vilar Sampaio, Juliano Coelho da Silveira, Felipe Perecin, Lawrence Charles Smith, Pablo Juan Ross, Flávio Vieira Meirelles

**Affiliations:** Department of Veterinary Medicine, Faculty of Animal Sciences and Food Engineering, University of Sao Paulo, Pirassununga, São Paulo, Brazil; Department of Animal Science, University of California, Davis, CA, USA; Centre de Recherche en Reproduction Animale (CRRA), Faculty of Veterinary Medicine, University of Montreal, Saint-Hyacinthe, Quebec, Canada

**Keywords:** Beta-hydroxybutyrate, ketone bodies, epigenetics, Histone Lysine β-hydroxybutyrylation, dairy cows, reproduction, cumulus cells, oocytes.

## Abstract

Besides their canonical roles as energy sources, short-chain fatty acids act as metabolic regulators of gene expression through the histone post-translational modifications. The ketone body β-hydroxybutyrate (BHB) was shown to cause a novel type of epigenetic modification, Histone Lysine β-hydroxybutyrylation (Kbhb), associated with genes upregulated in starvation-responsive metabolic pathways. Dairy cows increase BHB in early lactation and its effects on cellular epigenome are largely unknown. To unravel these effects, we sought and identified that Kbhb is present in bovine tissues in vivo and further confirmed that this epigenetic mark is responsive to BHB in bovine and human fibroblasts cultured in vitro in a dose-dependent manner. We also demonstrated that the maturation of cumulus-oocyte complexes with high concentrations of BHB did not affect the competence to complete meiotic maturation neither to develop until blastocyst stage. BHB treatment strongly induced H3K9bhb in cumulus cells, but this modification was only faintly detected in oocytes. Profiling the transcriptome in cumulus cells indicated that BHB treatment altered the expression of 345 genes. The down-regulated genes are mainly involved in glycolysis and ribosome assembly pathways, while the up-regulated genes are involved in mitochondrial metabolism and oocyte development. The specific genes and pathways altered by BHB treatment will provide entry points to carry out functional experiments aiming to mitigate problems and improve fertility in cattle suffering metabolic disorders. A key goal for future work will be to understand mechanistically how BHB transmits signals from the environment to affect cellular functions and the bovine epigenome.

**Summary sentence:** Beta-hydroxybutyrate induces Histone Lysine β-hydroxybutyrylation in fibroblasts and cumulus-oocyte complexes, it alters the transcriptome in cumulus cells, but does not affect oocyte’s competence to resume meiosis and develop until blastocyst stage.

## INTRODUCTION

Ruminant species represent a distinct class of animals with a specialized digestive organ, the rumen, that carries out the initial digestion of plant forage. The genome sequencing of domesticated ruminants has helped to elucidate particularities in the ruminant biology, such as the identification of genes involved in lipid metabolism showing particularities, such as of the roles of volatile fatty acids and ketone bodies that are different from non-ruminant animals [1, 2]. Ketone bodies, which include acetoacetate, acetone, and β-hydroxybutyrate, are generated in the liver mitochondria from the breakdown of fatty acids by β-oxidation and secreted in the bloodstream to serve as energy source.

Outstandingly, besides their roles as energy sources, recent findings using high-resolution mass spectrometry have shown a role of short-chain fatty acids as metabolic regulators of gene expression through the discovery of new histone post-translational modifications (PTMs) [3]. Metabolites from the cellular intermediary metabolism, such as succinate, propionate, crotonate, butyrate, among others, have been shown to cause a range of acylation reactions other than the classical and well-studied histone acetylation. These modifications of lysine residues within histones include Lys propionylation (Kpr), Lys butyrylation (Kbu), Lys 2-hydroxyisobutyrylation (Khib), Lys succinylation (Ksucc), Lys malonylation (Kma), Lys glutarylation (Kglu), and Lys crotonylation (Kcr). Similar to histone lysine acetylation (Kac), histone acylations are modulated by the cellular metabolism of cognate short-chain acyl-CoA species [4]. These modifications are implicated in the regulation of diverse physiological processes, such as signal-dependent gene activation, spermatogenesis, tissue injury, and metabolic stress [3].

In a landmark study, the ketone body β-hydroxybutyrate (BHB) was shown to cause a novel type of epigenetic modification, Histone Lysine β-hydroxybutyrylation (Kbhb), further expanding its epigenetic repertory [5]. In summary, Kbhb (1) is conserved and present in yeast, fly, mouse, and human cells; (2) is responsive and tuned to physiological (fasting, starvation) and pathological (diabetic ketoacidosis) variations of BHB concentrations; and (3) is correlated with gene transcription and marks a subset of genes upregulated in starvation-responsive metabolic pathways [5]. Thus, Kbhb is an epigenetic mark caused directly by BHB and can act as a link connecting the animal’s metabolic state with its chromatin.

Therefore, although usually regarded as a simple energy carrier, BHB can be ascribed to several surprising epigenetic functions for which the mechanisms remain elusive [6]. For instance, BHB is reported to protect against oxidative stress in the mouse kidney by inhibiting HDAC activity and thus enhancing histone Kac in promoters of genes driving anti-oxidative response [7]. Additionally, raising the levels of BHB through dietary intervention rescues hippocampal memory defects in Kabuki syndrome, a condition caused by mutations in enzymes involved in chromatin accessibility (i.e. KMT2D and KDM6A) [8]. Treatment of bovine cloned zygotes with BHB causes increased histone acetylation and blastocyst developmental rates, suggesting it may modulate chromatin accessibility and reprogramming pathways [9]. However, quantitative experiments (mass spectrometry and western blot) in both cultured cells and animals indicate that BHB has a much more profound impact on the levels of histone Kbhb than on acetylation [5]. Thus, the discovery of histone Kbhb and its prominent role in starvation-induced gene expression could help to elucidate metabolic problems that impact on the dairy industry, as well as on the reproduction of bovine and human beings.

The dairy industry faces a big problem related to nutritional imbalances due to a period of metabolic stress that cows undergo during the peri-calving period, the so-called negative energy balance (NEB) in the early weeks postpartum. NEB is characterized by increased circulating levels of nonesterified fatty acids (NEFAs) and BHB, which causes ketosis [10]. Epidemiological studies report that at least 50% of all dairy cows experience a temporary period of subclinical ketosis in the first month of lactation. Bovine ketosis is associated with reduction in milk production and correlated with health problems, e.g. metritis and abomasal displacement [11]. Moreover, metabolic stress in cows is associated with impaired fertility related to a difficulty to resume the ovarian activity and conceive in the first services [12, 13]. Since circulating BHB levels are mirrored in the follicular fluid of ovarian follicles [14], it is likely that BHB reaches the cumulus-oocyte-complex (COC) and could, thereby, jeopardize the developmental competence of the oocyte. Altogether, metabolic stress affects both the production and reproduction and is, therefore, a double-burden to farmers [15]. On the other hand, due to the health benefits demonstrated in clinical trials and laboratory studies with rodents, humans are adopting dietary regimens (e.g. ketogenic diet, intermittent fasting, time-restrict feeding) that lead to increases in BHB levels [16]. Moreover, human studies have used the ingestion of ketone esters to increase BHB levels and enhance metabolic performance in athletes [17]. Finally, other cataloged health benefits caused by BHB so far are diverse and range from improved memory, increased lifespan and reduction of hypertension [16]. Together, these observations show that there are many discrepancies regarding the role of BHB in humans versus cows, elucidating them can lead to new insights about the role of BHB in domestic animals and human reproduction.

Based on the data detailed above, there seems to be an intriguing and complex connection between bovine metabolism and epigenetics that can be further understood by studying Kbhb biology [4, 16]. To achieve this goal, we carried out a series of experiments aimed at: (1) determining whether histone lysine β-hydroxybutyrylation (Kbhb) is physiologically present in several tissues/organs in dairy cows; (2) examining whether supplementation with BHB *in vitro* increases the Kbhb in a dose-dependent manner in bovine and human fibroblast cultures; (3) exposing cumulus-oocyte complexes (COCs) during the *in vitro* maturation (IVM) to investigate whether BHB affects Kbhb levels in cumulus cells and oocytes; (4) determining whether the exposure of COCs to different concentrations of BHB during IVM alters the oocyte’s ability to complete meiotic maturation and develop to the blastocyst stage following parthenogenetic activation; and (5) characterizing the alterations in cumulus cell gene expression patterns (RNA-seq) in COCs exposed to the ketone body β-hydroxybutyrate during IVM.

## RESULTS AND DISCUSSION

### Identification of histone lysine β-hydroxybutyrylation in multiple bovine organs/tissues

During prolonged fasting or NEB period in cows, when glucose molecules are scarce, fatty acids replace glucose as the major energy source in the liver [18]. The organism initiates ketogenesis, where the acetyl-CoA is converted to ketone bodies such as BHB in the mitochondria of liver cells [19] that can either be exported to supply the organism with energy or be charged with free CoA to form β-hydroxybutyryl-CoA (bhb-CoA). Bhb-CoA is a high-energy donor for histone Lys β-hydroxybutyrylation (Kbhb) modification of which acyltransferase p300 catalyzes the transfer of BHB-CoA to histones [20] that mark and activate a subset of genes in response to starvation [5]. When the Kbhb modification is no longer necessary, it is removed by the histone deacetylases HDAC1 and HDAC2 [20]. This model is depicted below (**Figure 1A**) and explains how fluctuations in the levels of BHB derived by nutritional cues can increase Kbhb, tag histones, and trigger a gene expression response [3].

**Figure 1.**
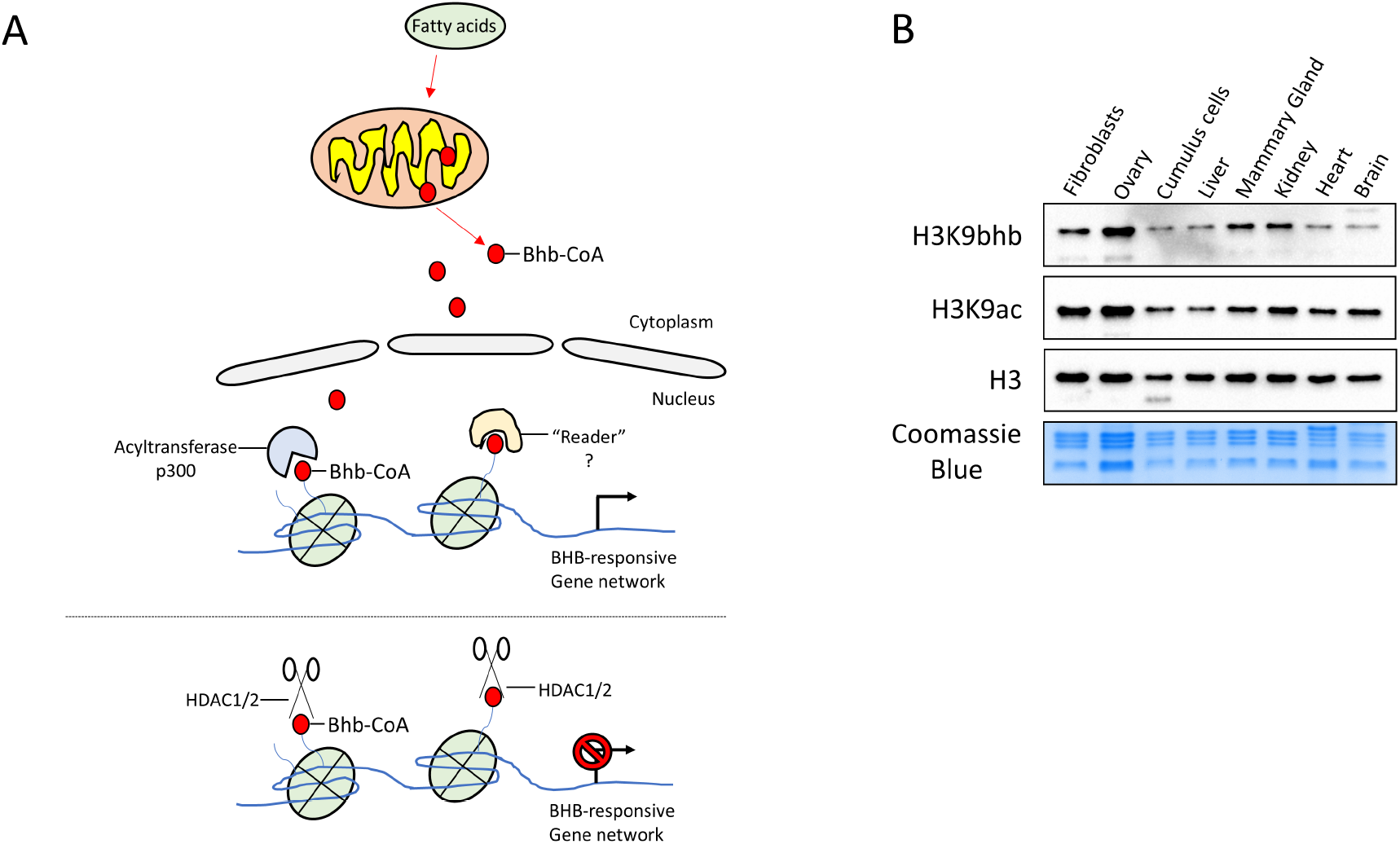
**A-** Illustration depicting the postulated mechanisms by which Histone Lysine β-hydroxybutyrylation is produced, deposited in histones, and read or removed in a given cell. During early lactation in cows or prolonged fasting in mammals, when glucose molecules are scarce, fatty acids replace glucose as the major energy source in the liver. In ketogenesis, acetyl-CoA is converted to ketone bodies such as β-hydroxybutyrate (red circles) in the mitochondria of liver cells and exported to supply peripheral tissues. Alternatively, β-Hydroxybutyrate can be charged with free CoA to form β-hydroxybutyryl-CoA (bhb-CoA). The high-energy donor for the histone Lys β-hydroxybutyrylation (Kbhb) modification can be transferred by the histone acetyltransferase p300 to histones. Kbhb marks a class of genes that are activated in response to starvation. When the animal returns to a well-fed state, Kbhb is removed by Histone desacetylases HDAC1 and HDAC2 and the starvation-responsive genes are silenced. The question mark on the “reader” protein denotes it has not been identified so far. **B-** Coomassie blue stained gel showing enrichment for histone proteins and immunoblots for H3, H3K9ac and H3K9bhb in dairy cow cells and tissues. The western blot images were cropped for illustrative purposes. The full gels/blots are presented in Supplemental Figure 1.

Histone Lysine β-hydroxybutyrylation (Kbhb) is a novel epigenetic post-translational modification that was identified in the liver of mice subjected to fasting or streptozotocin-induced diabetic ketoacidosis [5]. Moreover, Kbhb was identified in the brains of depressive mice, indicating a role in modulating mood and suggesting it is physiologically necessary even at basal levels [21]. Dairy cows commonly undergo ketosis episodes during lactation and so far, the post-translational modification (PTM) Kbhb has not been demonstrated in cattle. To investigate whether Kbhb is present *in vivo* in cattle, we collected samples from dairy cow ovary, cumulus cells, liver, mammary gland, kidney, heart, brain, and from a fibroblast cell culture. We acid-extracted histones and run an SDS-PAGE gel to confirm histone enrichment by Coomassie blue staining (**Figure 1B**). Next, we carried out western blot to identify Kbhb utilizing an antibody for the H3K9bhb residue. The choice for the H3K9bhb antibody was due to its robust response to starvation in mouse liver, and specificity by dot blot assays and competition experiments [5]. As a result, we were able to confirm that Kbhb (hereafter we will use H3K9bhb or Kbhb interchangeably throughout the manuscript) is present in all organs and cells investigated (**Figure 1B**). As controls, we used immunoblot for total H3 and H3K9ac, another PTM of the same lysine residue, and, as expected, found that both were present in all samples analyzed.

Regarding physiological relevance, mice starved for 48 hours showed an increase in liver histone Kbhb that was highly correlated with the transcription of genes involved in a starvation-responsive pathway [5]. Kbhb is reduced in the brain of two mouse models of depressive behavior, i.e. dexamethasone and spatial restraint stress models. BHB increment using a ketogenic diet or through exogenous BHB supplementation led to the reexpression of the neurotrophic factor BDNF, increased H3K9bhb and showed antidepressant effect [21]. During fasting, levels of histone acetylation increase in the kidney, are maintained in the liver and decrease in the heart, indicating different responses in each organ [19]. These findings suggest that the inhibition of HDACs by BHB may be context-dependent and also that each organ possesses a different acetylome even in conditions affecting the animal systemically. In relation to the organ-specific patterns of Kbhb, although our western-blot approach can only provide a semi-quantitative exploratory assessment, we observed a strong signal in the ovary. Previous reports have found dramatic changes in ovarian cells during the different stages of the estrous cycle in dairy cattle, suggesting a high metabolic rate in this organ. Indeed, cows have been shown to have a significantly higher uptake of BHB in their ovaries compared with NEFAs [22], suggesting that cattle may utilize BHB as a source of energy during periods of nutritional imbalance. Beside its role as energy source, we speculate that BHB might modulate the activity of luteal or stromal cells through Kbhb and provide a link between nutrition-ovarian function-reproduction in cows. Isotope-labeled carbon studies demonstrated that the brain and the mammary gland are major sites of ketone body incorporation into lipids [23]. Because elevated BHB levels are correlated with reduction in milk production and altered lipid composition, BHB may have another fate in mammary gland and epigenetically activate enzymes responsible for lipid metabolism. Supporting this notion, β-hydroxybutyrate generated immediately after birth activated postnatal gene expression in the mouse neonate liver by epigenetic mechanisms [24]. Since the activated genes were mostly involved in milk lipid catabolism, Kbhb-associated gene expression may contribute additionally to this task. Future studies will require the combination of techniques to map the distribution of Kbhb in a genome-wide manner, i.e. by ChIP-seq or Cut&Tag, with RNA-seq in each organ of interest and under different metabolic states (e.g. NEB cows) to unravel which regions are enriched in Kbhb and the genes that are activated in each organ to provide clues about the function of Kbhb-associated epigenetic modifications.

### Supplementation of bovine fibroblasts with BHB increases H3K9bhb in a dose-dependent manner

Post-translational modifications (PTMs) to histones depend on small-molecule metabolites such as acetyl-CoA, butyrate, succinate, β-hydroxybutyrate, among others [25]. Thus, due to this dependency on such metabolites, PTM can relay information about cellular metabolic state to the genome for the activation or repression of particular sets of genes [3]. The levels of a given histone acylation are directly related to the cellular concentration of its specific acyl-CoA. In mice, Kbhb increases in the liver by up to 40-fold in relation to the increment in BHB levels caused by physiological (e.g., fasting) or pathological conditions (e.g., diabetic ketoacidosis). Treatment of cells with heavy carbon-labeled BHB leads to the labeling of histone proteins with a heavy Bhb-Coa, suggesting that BHB is converted into its cognate acyl-CoA, which is then used directly as a cofactor to catalyze Kbhb. Such data was further confirmed in experiments by western blot in which Kbhb increased in HEK293 cells proportionally to treatments with exogenous BHB [5].

That said, high-yielding dairy cows face metabolic disturbances in which BHB levels can rise from basal/physiological levels (i.e., ∼0.5 mM) to as high as 6 mM during severe clinical ketosis [26]. To verify whether Kbhb increases proportionally to BHB levels in cattle, we used skin fibroblasts as a tractable platform for sodium β-hydroxybutyrate treatment. We chose the concentrations of 0 mM (control), 2 mM, 4 mM, and 6 mM of BHB with the rationale that these concentrations are in the range (from basal levels to the highest found in cattle) measured in animals suffering from subclinical, clinical and severe clinical ketosis [27]. Additionally, we used the 5 mM Sodium Butyrate (hereafter named NaBu), a canonical inhibitor of histone deacetylases [28], as a positive control to measure histone acetylation. After a 24 h treatment, we measured the intensity of Kbhb (antibody against the H3K9bhb, residue critical for transcriptional regulation) utilizing immunostaining followed by confocal microscopy. Results indicate that supplementation of cell culture with sodium β-hydroxybutyrate increases H3K9bhb in fibroblasts in a dose-dependent manner (p<0.0001). When compared with controls, cells treated with 2 mM BHB possess ∼ 2.56-fold more H3K9bhb staining compared to untreated cells, while cells treated with 4 and 6 mM BHB show 3.91 and 5.22-fold increases, respectively (**Figure 2 A-F**). To further confirm our data using confocal-based immunofluorescence intensities, we performed an experiment using 3 independent biological replicates of fibroblasts, acid-extracted histones and carried out western blot. As expected, we observed an increase of 2.27, 4.75, and 6.62-fold change in cells treated with 2, 4, and 6 mM BHB compared to untreated controls **(Figure 2 G-H), p=0.0002**. These data demonstrate that Kbhb is highly responsive to BHB exposure and increases Kbhb levels significantly after BHB supplementation in a dose-dependent manner, suggesting that alterations in the circulating levels of BHB observed during ketosis could influence the bovine epigenome by activating genes to respond to the metabolic stress. Regarding histone acetylation, BHB supplementation did not affect H3K9ac in cultured cells. As expected, supplementation with NaBu increased histone acetylation (∼2-fold) compared with all the other treatments (Supplemental Figure 2). These observations support previous findings where the increment in acetylation is minimal or absent, probably due to the use of culture media with artificially high glucose concentrations and/or the predominant effect of BHB on Kbhb over Kac [3,5,9,21].

**Figure 2.**
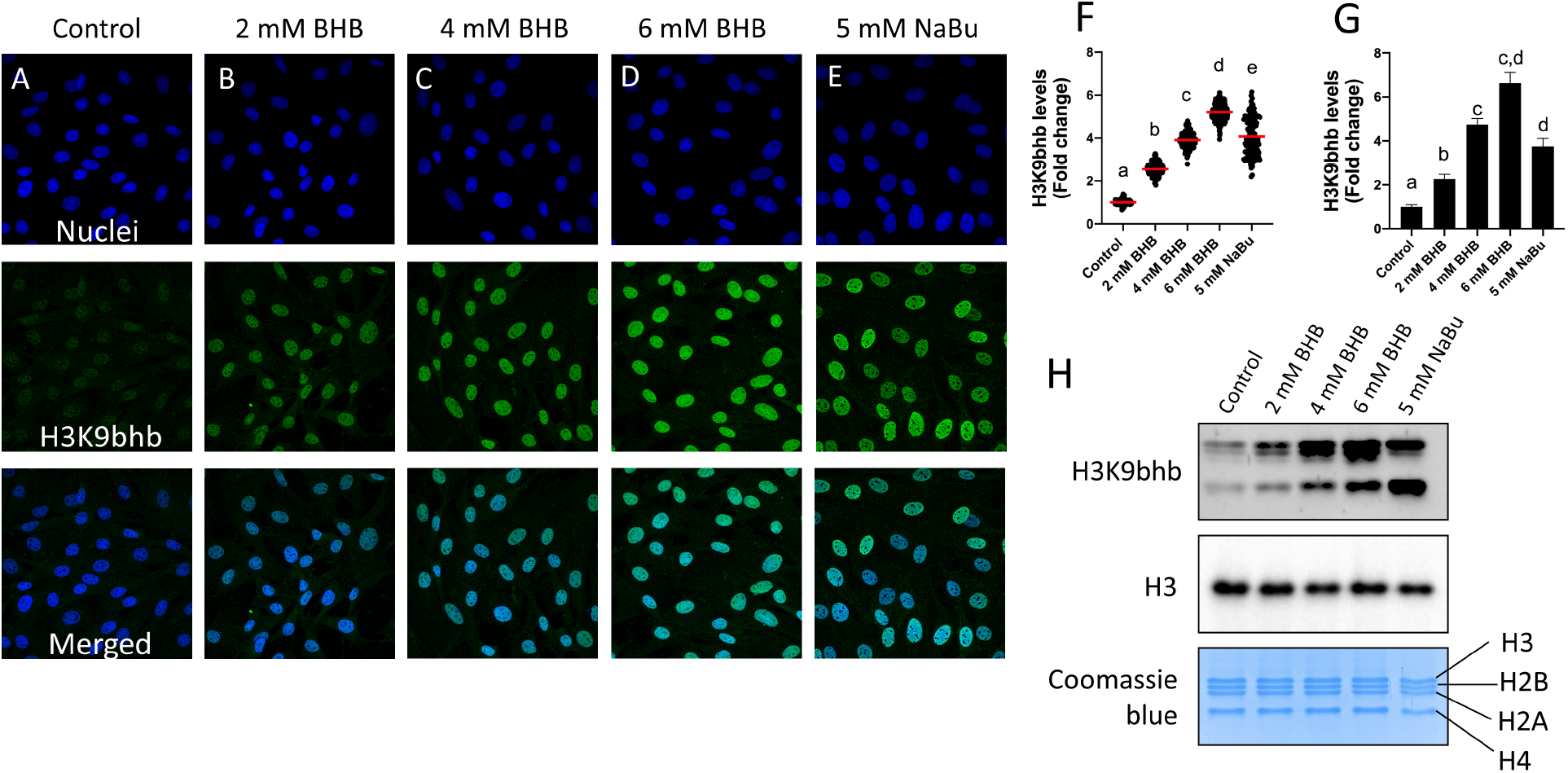
Immunostaining of H3K9bhb in bovine fibroblasts cultured for 24 h in culture medium supplemented with **(A)** 0, **(B)** 2, **(C)** 4, **(D)** 6 mM BHB, and **(E)** 5 mM NaBu. **(F)** Scatter plot showing H3K9bhb fluorescence levels in fold change and the mean (red dash) in nuclei (n>120) of fibroblasts measured by confocal microscopy. Treated group values are represented as fold change compared to the control group. Different letters indicate statistical differences between groups as measured by Tukey’s test (p<0.05). **(G)** Bar graph shows the H3K9bhb levels in fold change (± sem) measured by Western blot (WB). **(H)** Representative gel stained with coomassie blue and immunoblots for H3K9bhb and total H3 from bovine fibroblasts cultured as described above. The western blot images were cropped for illustrative purposes. The full gel/blots are presented in Supplemental Figure 3. Nuclei were stained with Hoechst 33342 and H3K9bhb with Alexa Fluor 488 dye. Pictures were taken at 63x magnification.

Curiously, we observed an increment (∼4.07 fold confocal and ∼3.76 WB) of Kbhb in fibroblasts treated with NaBu when compared to untreated controls (**Figure 2E-H**). A recent report utilizing human cells also demonstrated that NaBu supplementation increased the levels of Kbhb, suggesting a conserved effect among mammals [29]. Supporting our findings, it was shown that Kbhb is removed by histone deacetylases and treatment with broad class I and II histone deacetylases inhibitors (e.g. TSA and NaBu) caused Kbhb accumulation in cultured cells [20]. Alternatively, since butyrate can be converted to β-hydroxybutyrate in the bovine ruminal epithelium [30] and supposing the same reaction occurs to in vitro cultured fibroblasts, the supplementation of cells with butyrate may generate BHB that is later charged to BHB-CoA and tag histones.

The core histone proteins H3, H2B, H2A, and H4 migrate in SDS-PAGE generating a similar banding pattern for most mammalian species [31]. As observed in the Coomassie blue stained gel (Figure 2H), our electrophoresis separated the 4 histones in the expected pattern. The H3 migrates slower, followed by H2B, H2A and H4 (Figure 2H). Another observation from our western blots is that the antibody becomes less specific and recognizes other β-hydroxybutyrylated sites on histones H2A, H2B and H4 proportionally to the BHB concentrations (Figure 2H). Several studies have observed few problems regarding the behavior of histone PTM antibodies, including off-target recognition, strong influence by neighboring PTMs, and an inability to distinguish the modification state on a particular residue, such as mono-, di-, or trimethyl lysine [32]. Another problem is the structural similarity presented by some histone acylation [3]. However, extensive validations for this antibody have been performed by dot blot and competitive assays showing it does not react against acetylated, butyrylated, propionylated, among other acylations [5]. Recent work has demonstrated that the radical responsible for Kbhb, β-hydroxybutyryl-CoA is highly reactive and can tag histones by nucleophilic attack [33]. Our data indicates that exogenous supplementation of BHB dramatically increases Kbhb levels, and that in this condition the antibody does not properly distinguish the specific residue (H3K9bhb) for which it was raised. Nonetheless, such slight cross-reactivity does not affect the general conclusion that supplementation with exogenous BHB increases Kbhb in a dose-dependent manner in fibroblasts. Further identification and characterization of Kbhb readers or effectors proteins need to be performed to fully decipher the Kbhb code in animals [19, 34].

### Supplementation of human dermal fibroblasts with BHB increases H3K9bhb in a dose-dependent manner

Humans are adopting several practices (e.g. ingestion of ketone ester drinks) and dietary regimens (e.g. ketogenic diet, intermittent fasting, time-restricted feeding) aiming to improve wellness, athletic performance, and health [17,35,36]. In humans, serum levels of BHB are usually in the low micromolar range (50 μmol/L in the normal fed state), but can begin to rise in some physiological and pathological conditions. For instance, BHB levels reach ∼1–2 mM after 2 days of fasting or 90 min of intense exercise, and 6–8 mM with prolonged starvation (∼4 weeks) [37]. Outstandingly, in a life-threatening condition known as diabetic ketoacidosis, BHB levels can reach ∼20 mM [38]. Attaining a ketogenic diet that is almost devoid of carbohydrates, consistent levels above 2 mM are also reached. The ketone body BHB has been demonstrated to be a key molecule mediating the beneficial effects of diets or interventions [16]. Thus, we decided to investigate the effects of BHB in human fibroblast cultures and compare them to our bovine findings.

For such, we treated human dermal fibroblast (hDF) with control, 2 mM, 6 mM and 20 mM of BHB for 24 h. Similarly to bovine fibroblasts, hDF significantly increased (p<0.0001) Kbhb levels in a dose-dependent manner (**Figure 3A-E**). Cells treated with 2 mM increased ∼2.21-fold in relation to untreated controls (**Figure 3F-G**), while 6mM BHB caused a ∼5.97-fold increase in Kbhb (**Figure 3F-G**) as measured by western blot. Regarding the pathological concentrations of 20 mM BHB, Kbhb levels were dramatically increased by 10.22-fold (**Figure 3F-G**). As observed by confocal microscopy, image analysis of hDF cells treated with 20 mM (**Figure 3D**) show saturation of Kbhb fluorescence levels with increases of over 50-fold in comparison to controls (**Figure 3E**). It is noteworthy that the laser power of the confocal microscope was adjusted to avoid saturation in physiological concentrations (up to 6 mM). Together, these data indicate that Kbhb levels are highly responsive and dependent on BHB concentrations in different mammals. Regarding histone acetylation, previous studies with HEK 293 cells [5] and primary neurons [21], have demonstrated that cultures in the presence of beta-hydroxybutyrate up to 20 mmol/L only marginally increases histone acetylation (Kac). We further confirmed this finding in hDF, demonstrating that BHB has a more pronounced effect on Kbhb than Kac (Supplemental Figure 4).

**Figure 3.**
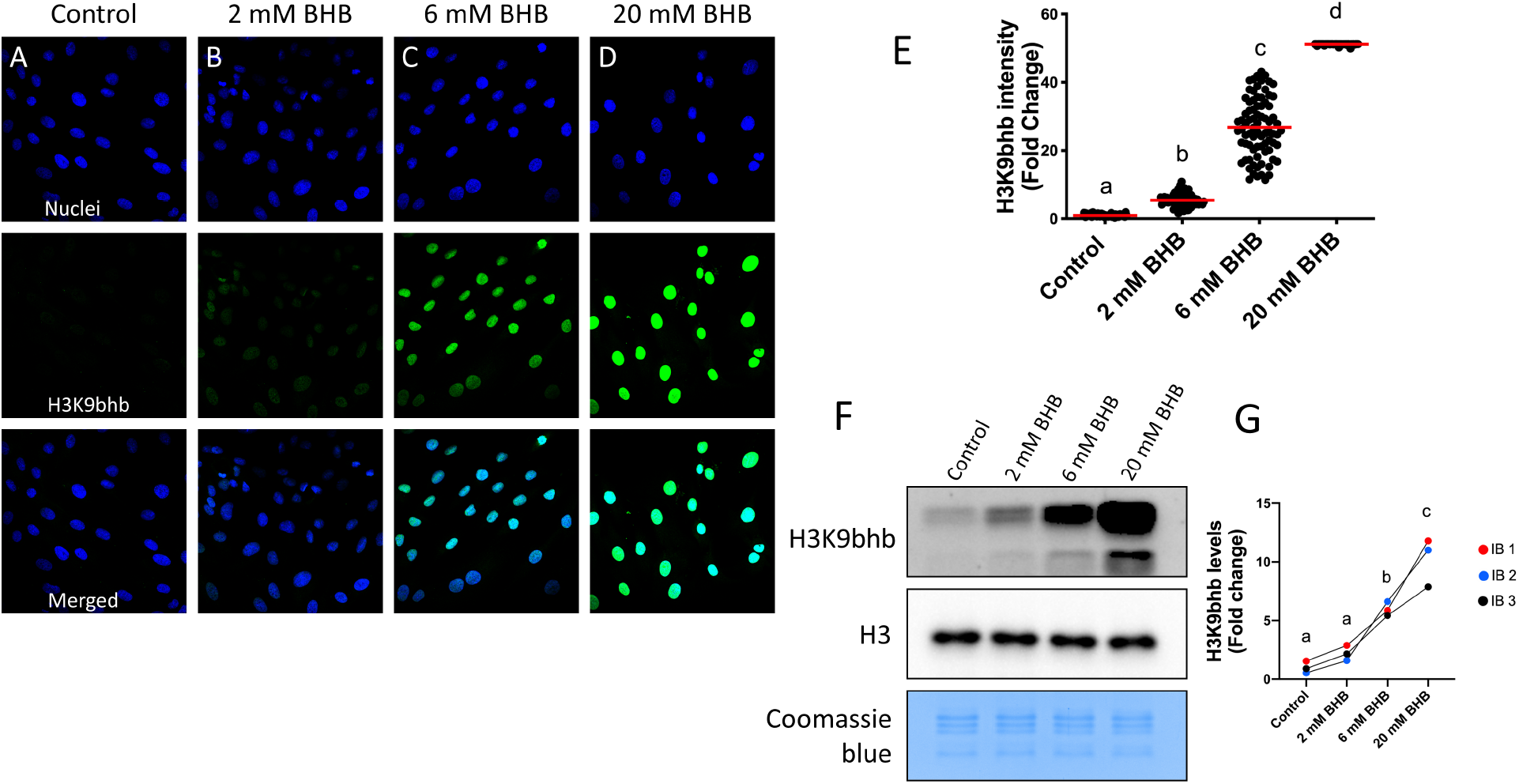
Immunostaining for H3K9bhb in human dermal fibroblasts (hDF) cultured for 24 h in medium supplemented with **(A)** 0, **(B)** 2 **(C)** 6, and **(D)** 20 mM BHB. **(E)** Scatter plot showing H3K9bhb fluorescence levels in fold change and the mean (red dash) in nuclei (n>65) of hDF measured by confocal microscopy. Different letters indicate statistical differences (p<0.0001) using Kruskal-Wallis’ test. **(F)** Representative gel stained with coomassie blue and immunoblots for H3K9bhb and total H3 from hDF cultured as described above. The western blot images were cropped for illustrative purposes. The full gel/blots are presented in Supplemental Figure 5. **(G)** Graph shows the intensities of H3K9bhb bands in fold change from the immunoblots (IB). Different letters indicate statistical differences (p<0.0001) using Tukey’s test. Each color represents an experimental replicate. Nuclei were stained with Hoechst 33342 and H3K9bhb with Alexa Fluor 488 dye. Pictures were taken at 63x magnification.

The skin has been disregarded in the vast majority of metabolic studies that focus mostly on major organs, such as the liver and the intestine. However, skin is the largest mammalian organ and acts as a primary barrier against heat loss, dehydration, mechanical trauma, and microbial insults [39]. As a consequence, skin is expected to be important in adaptations to stimuli such as dietary interventions with an effect on aging. Indeed, a previous report has indicated a beneficial effects of dietary interventions such as caloric restriction on metabolic adaptations and regeneration in skin [40]. BHB is known to participate in stem cell proliferation and regeneration in intestines [41]. Since our treatment demonstrated that human and bovine fibroblasts are highly responsive to BHB supplementation, it is expected to act as a nexus connecting dietary interventions with skin renewal. The implications of the epigenetic alterations caused by BHB on fibroblasts warrant further investigation.

### Turnover of Kbhb in bovine fibroblasts happens quickly after BHB withdrawal

Epigenetic marks on chromatin are dynamically changing in the cell nuclei to allow a rapid response to external stimuli and to regulate gene expression. Histone acylations are regulated by a dynamic balance between acyltransferases and deacylases. Among the epigenetic marks, acetylated histones are turned over very quickly, i.e. within minutes with a half-life of acetylation of about 2–3min. On the other hand, histone methylation displays a much slower turnover rate with a half-life of about 0.3–4 days [42]. Surprisingly, there is no information about the turnover rate of Kbhb. Therefore, studies on Kbhb turnover is important to understand the rate at which Kbhb deposition resulting from an episode of fasting or ketosis is withdrawn once the animal returns to a well-fed state. For instance, if such a modification lingers in the epigenome, it could physically block the deposition of other epigenetic modifications and affect subsequent developmental processes, such as cellular differentiation, oocyte growth and embryonic development [25].

To investigate the decay of Kbhb, we treated bovine fibroblasts with 6 mM BHB and carried out immunostaining at different time points after removal of the BHB treatment. As observed in the **Figure 4A-B** **and 4G**, treatment with BHB causes a 6-fold increment in the intensity of Kbhb fluorescence in fibroblasts (p<0.0001). During BHB withdrawal, Kbhb intensity levels in fibroblast nuclei drop by∼11% after 1 h (5.4-fold; **Figure 4C** and **4G**), by ∼28 % after 2 h (4.4-fold; **Figure 4D** and **4G**), and by roughly 50% after 4 h (3.1-fold; **Figure 4E** and **4G**) after withdrawing Kbhb from the culture medium. These data demonstrate that, once the exposure to BHB is withdrawn from the cultured fibroblast, Kbhb is rapidly removed from the chromatin, simulating the end of a fasting episode and the return to a “well-fed state”. Evaluation at 24 h after BHB removal indicated a ∼75% drop (1.48-fold, **Figure 4F** and **4G**) in Kbhb fluorescence levels. Nonetheless, in spite of the sharp decline in Kbhb levels even after 24 h of BHB removal, the cells still showed elevated Kbhb levels (∼1.48-fold) in comparison to untreated controls (**Figure 4G**), suggesting that either some sites on the chromatin are resistant to BHB withdrawal or associated with genes involved in late-response to nutrient deprivation/starvation.

**Figure 4.**
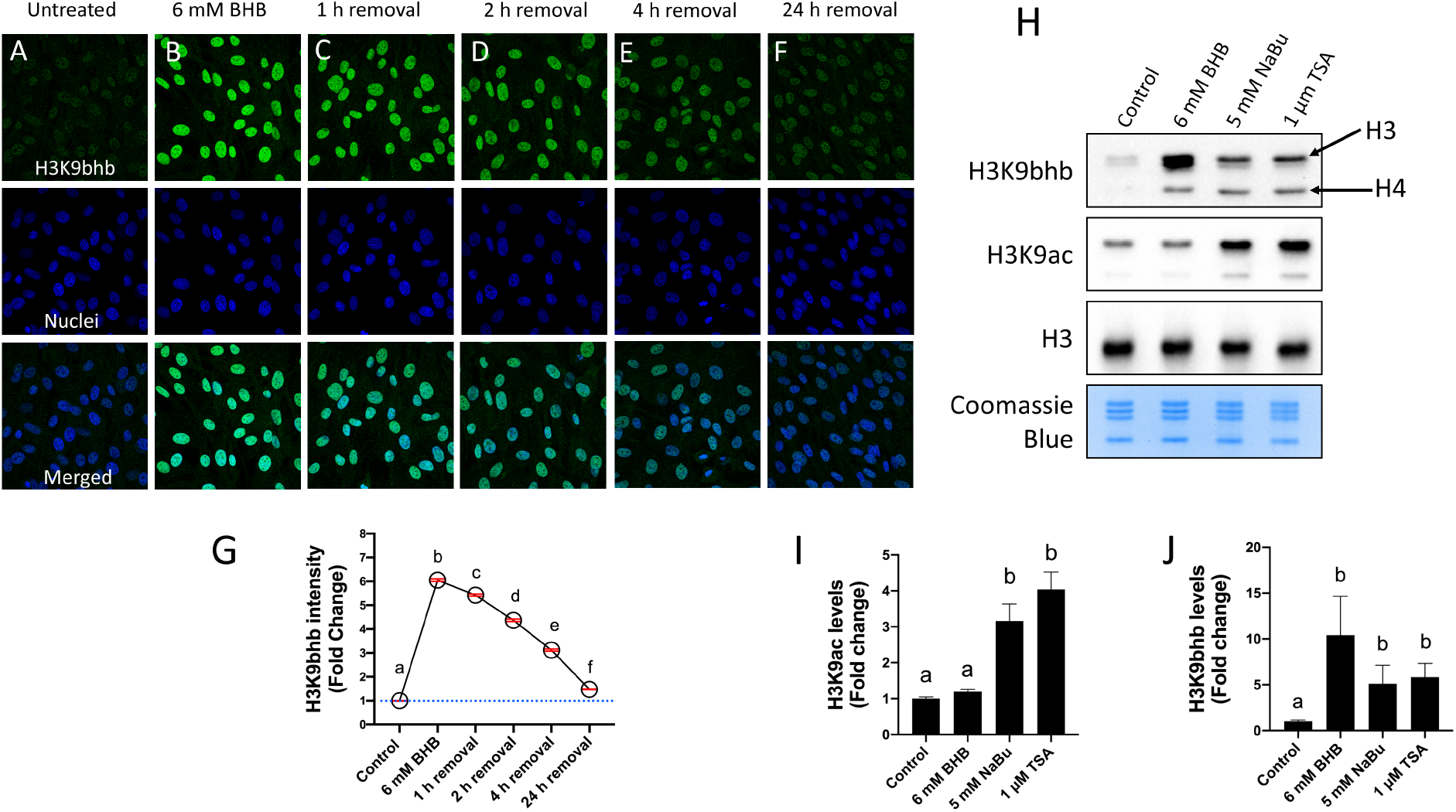
Immunostaining for H3K9bhb in bovine fibroblasts cultured for 24 h in culture medium supplemented with **(A)** 0 and **(B)** 6 mM BHB. Additionally, BHB culture medium was replaced by control medium and cultured for **(C)** 1, **(D)** 2, **(E)** 4 and **(F)** 24 h after BHB removal. **(G)** Graph shows the H3K9bhb intensity levels (fold change ± sem) in fibroblasts measured by confocal microscopy. The blue dashed line shows the average fluorescence levels in untreated cells. In red are the standard error of the mean. **(H)** Representative gel stained with coomassie blue and immunoblots for H3K9bhb, H3K9ac, and total H3 from bovine fibroblasts from control and treated groups exposed for 24 h to 6 mM BHB, 5 mM NaBu and 1 µM TSA. The western blot images were cropped for illustrative purposes. The full gel/blots are presented in Supplemental Figure 6. The black arrows are pointing the antibody recognizing the H3 and the H4 based on the expected electrophoretic mobility **(I)** Bar graph shows the H3K9ac levels (fold change ± sem) in fibroblasts as measured by WB. **(J)** Bar graph shows the H3K9bhb levels (fold change ± sem) in fibroblasts as measured by WB. Different letters indicate statistically significant difference (p<0.05) among groups as assessed by Tukey’s test. Nuclei were stained with Hoechst 33342 and H3K9bhb with Alexa Fluor 488 dye. Pictures were taken at 63x magnification.

The rapid removal of most of the Kbhb signal suggests that an active mechanism may be operating to remove this epigenetic mark in cattle. Histone deacetylases were demonstrated to remove Kbhb from core histones in vitro from HEK293 cells [20]. Since our previous findings (**Figure 2E-H**) showed an increase in Kbhb levels after exposure to NaBu, presumably by blocking HDAC activity, these results further support the notion that HDACs are responsible removing Kbhb. As mentioned previously, another possibility is that NaBu is converted to BHB in the cell culture and, thus, could donate the radical for Kbhb deposition. To test the hypothesis that the accumulation of Kbhb occurs by HDAC inhibition, we examined the effect of another chemically unrelated and broad histone deacylase inhibitor (HDACi), i.e. the fungal compound Trichostatin A (TSA) [43]. To first validate that NaBu and TSA inhibited histone deacetylases, we blotted for H3K9ac, a marker for histone acetylation. As expected, NaBu increased ∼3.16-fold (p= 0.0090) the global levels of histone acetylation and TSA, a more potent HDACi, ∼4.04-fold, p=0.0011 (**Figure 4H-I**). Surprisingly, treatments with 6 mM BHB (positive control), NaBu and TSA all led to an increase in Kbhb, demonstrating that a broad range of HDAC inhibitors causes Kbhb accumulation (**Figure 4H and 4J**). BHB treatment increased ∼10.41-fold the levels of H3K9bhb, while NaBu and TSA increased over 5-fold (**Figure 4J**), p<0.05. Therefore, accumulation of Kbhb after TSA treatment, a molecule chemically unrelated to BHB, further supports the notion that H3K9bhb is removed by HDACs in cattle.

Sirtuins also have risen as potential candidates to act as erasers. Although sirtuins are cataloged as NAD^+^-dependent histone desacetylases, growing evidence suggests that sirtuin enzymes have an expanded repertoire of deacylation activities [3]. A systematic survey of the activity of the seven known mammalian sirtuins against synthetic peptides containing several histone acylations showed that they have depropionylase, debutyrylase, decrotonylase activities, among others [3]. The wide diversity of lysine acylations removed by the various mammalian sirtuins hints that hydroxybutyrylation might be also removed by a specific sirtuin. Indeed, previous studies have demonstrated that SIRT 1 and SIRT 2 have de–β-hydroxybutyrylation activity [20]. In another study, Sirtuin 3, a mitochondrial deacetylase, was recently demonstrated to be “moonlighting” in the cellular nucleus and removes Kbhb in some specific residues, included H3K9bhb used herein [44]. However, Sirtuin 3 showed strong selectivity for the S-form of BHB while animals usually generate the R-form in physiological conditions [19].

Regarding the deposition of Kbhb, based on mass spectrometry data, β-hydroxybutyrylation occurs to a greater extent through non-enzymatic mechanisms in both histones H3 and H4 [33]. Curiously, based on our immunoblots, cells treated with NaBu and TSA tend to accumulate more signal at the position corresponding to the Histone H3 and H4 (see black arrows in **Figure 4H**; based on the electrophoretic mobility). This fact is curious and supports the idea that using these broad HDAC inhibitors favors the equilibrium towards the accumulation of Kbhb that happens by two postulated mechanisms. First, the nucleophilic attack occurs on histones and the inhibition by NaBu and TSA hampers Kbhb removal, leading to accumulation. Second, the acyltransferase p300 has been shown to have histone Kbhb transferase activity in cells and to stimulate transcription in vitro, demonstrating that, similar to Kac, Kbhb is controlled by enzymes with opposing activities (i.e. p300 and HDACs) [20]. All these discrepancies regarding the Kbhb metabolism need to be addressed in the future to further elucidate the Kbhb code in mammals. Moreover, key regulatory elements of Kbhb pathway, including the “readers” remain largely unknown, thereby hampering its biological characterization [19, 34].

As possible implications of our findings, butyrate is an important food source for colonocytes, the epithelial cells that line the colon, that use it to fuel β-oxidation for energy production and also to inhibit the growth of cancerous, but not noncancerous, colonocytes [45]. A pathway connecting BHB, inhibition of HDAC, and Notch signaling has been demonstrated to regenerate the intestinal epithelium [41]. Since butyrate can inhibit some deacetylases responsible for BHB removal, it could provide a new layer of mechanisms by which rich fiber diets enhance butyrate production by colonic bacteria, causing inhibition of Kbhb removal and epigenetics effects similar to BHB coming from other sources (e.g. ketogenic diet) in intestines.

### Exposure of cumulus-oocyte complexes to BHB during initial stages of in vitro maturation increases the H3K9bhb levels in cumulus cells

After identifying and characterizing the effect of BHB in organs and cultured cells, we carried out a series of experiments to investigate the effect of BHB in bovine cumulus-oocyte complexes (COCs) during in vitro maturation to further elucidate the role played by BHB in a context of bovine reproduction.

Cumulus cell–oocyte communication is an essential process to establish oocyte competence involving a bidirectional transfer of ions and small molecules. Cumulus cells utilize transzonal projections (TZPs) and gap junctions to transfer molecular cargoes that are critical for regulating the process of oocyte maturation [46]. On the other hand, oocytes use structures named oocyte-derived mushroom-like microvilli [47] and paracrine signaling to regulate much of the flow of ions and molecules towards the cumulus [48]. In the past few years, a less well-characterized and new potential mechanism has been identified that involves the exosomal transfer of noncoding RNAs from cumulus to oocytes [49, 50]. Indeed, by studying nascent RNA with confocal and transmission electron microscopy in combination with transcript detection, it was showed that the somatic cells surrounding the fully-grown bovine oocyte contribute to the maternal reserves by actively transferring large cargoes, including mRNA and long noncoding RNA [51].

Due to the BHB epigenetic role in activating or being associated with gene networks involved in starvation response [5], we sought to investigate the effect of COCs exposure to BHB on Kbhb levels during the firsts hours of in vitro maturation. We matured COCs with 2 mM BHB for 4 h, followed by an increase to 4mM BHB for an extra period of 4 h (by adding 2 mM of BHB solution directly in the oocyte’s maturation dish, see materials and methods section), totalizing 8 h BHB treatment during the initial stages of in vitro maturation. The rationale for this short period of exposure was: (1) To examine whether and how soon BHB treatment induces the appearance of Kbhb; (2) Using an in vitro model, to mimic a hypothetical situation where a cow is abruptly exposed to BHB during severe ketosis (**Figure 5 A**); (3) Since transzonal projections are completely functional during the firsts 8 h of maturation, they potentially are able to transfer functional molecules, such as coding and noncoding RNAs, from the cumulus cells to the oocyte [52]. After BHB treatment, western blot results indicated that Kbhb levels increased ∼2-fold in treated COCs compared to untreated controls (**Figure 5B-C**). Together, these data indicate that a short exposure of COCs to BHB triggers Kbhb accumulation in cumulus cells.

**Figure 5.**
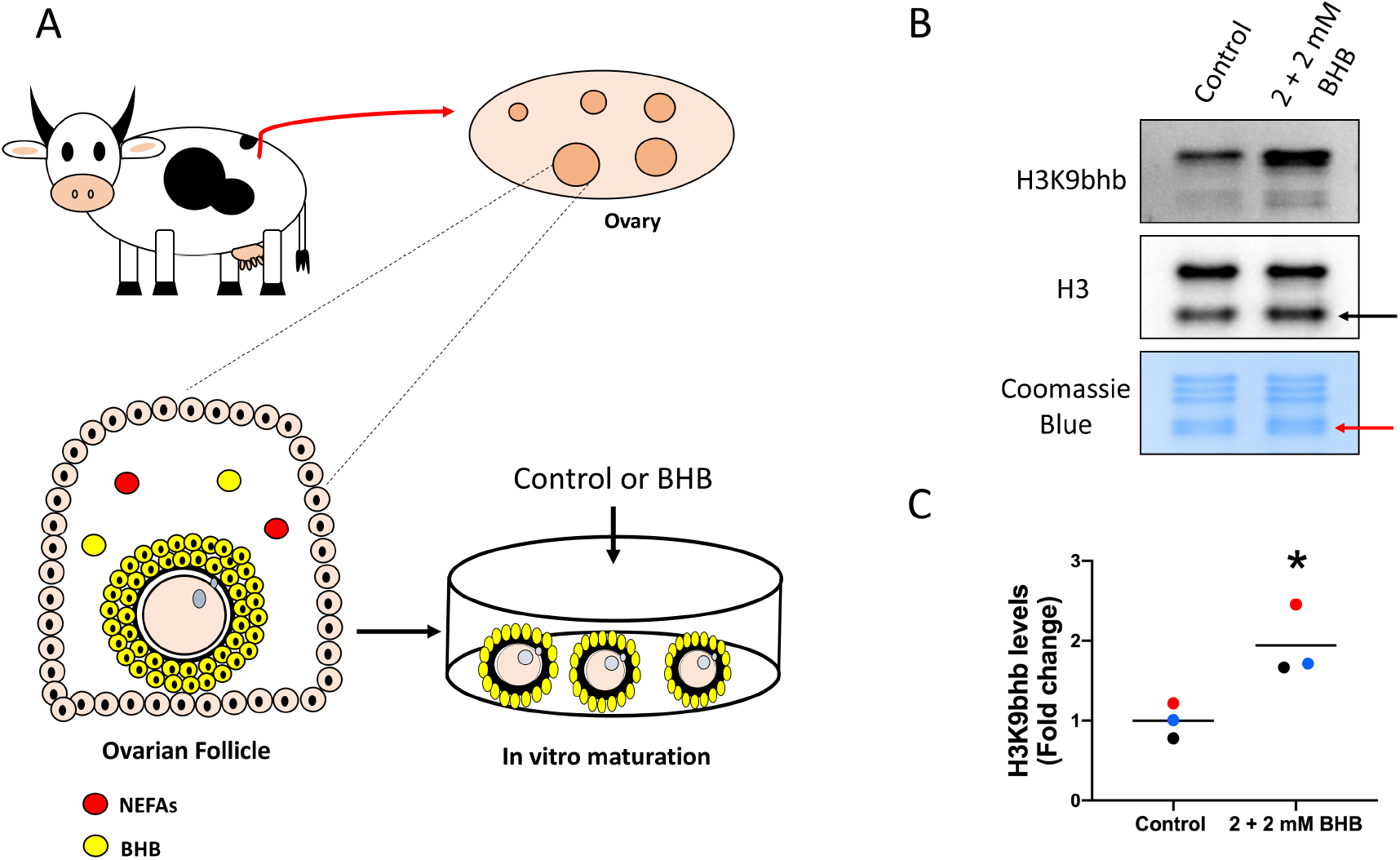
**(A)** Illustration depicting the microenvironment of an ovarian follicle, in which metabolites such as non-esterified fatty acids (NEFAs) and BHB are present in vivo at similar concentrations to the plasma. We aimed to mimic in vivo conditions by maturing COCs in vitro with high concentrations of BHB to investigate the effects on COCs epigenome and development. **(B)** Representative gel stained with coomassie blue and immunoblots for H3K9bhb and total H3 from cumulus cells cultured for 8 h in IVM medium supplemented with 2 to 4 mM BHB or not (Control). The western blot images were cropped for illustrative purposes. The full gel/blots are presented in Supplemental Figure 7. The black arrow is pointing an extra band observed in histones extracted from cumulus cells. The red arrow is pointing the region where the H4 is commonly observed, but showing a second band not observed in other tissues extracted using the same protocol. **(C)** Graph shows the H3K9bhb levels in fold change in cumulus cells measured by WB. Asterisk (*) indicates statistical difference (p=0.0295) using Student’s t-test. BHB treated groups are represented in fold change in relation to controls. Each color represents specific experimental replicates and the black horizontal bars represent the group average.

As an evolving field, virtually all metabolites from the intermediary metabolism have been shown to affect the cellular epigenome [53]. Moreover, recent findings have indicated a high correlation of Kbhb with gene expression, indicating a causative role in metabolite-induced alterations to gene expression [54]. Thus, it will be enlightening to investigate whether cumulus cells are able to ship metabolite-stimulated cargoes to oocytes during the final period of oocyte maturation. Such findings could be evolutionarily important because, after undergoing the whole period of oocyte growth, oocytes could still receive dietary-stimulated RNA molecules (or others such as lipids and metabolites) informing about the prevailing nutritional condition faced by the animal close to the periconceptional period [55]. However, both the consequence and impact of the transfer of such large cargoes via exosomal communication pathways remain poorly understood in mammalian gametes [56].

### BHB exposure of COCs during the entire in vitro maturation period does not affect the maturation rates nor the developmental capacity of oocytes

A healthy intrafollicular environment is necessary during the acquisition of developmental competence of oocytes. An oocyte is considered competent when it is able to resume meiosis, undergo fertilization, embryogenesis, and generate a viable offspring [57]. The ovarian follicle becomes highly vascularized during the advanced antral phase, and the follicular fluid (FF) mirrors some components from the systemic environment [14]. Although such components (e.g. BHB, NEFAs, glucose) nurture the developing COCs in preparation for ovulation, harmful compounds can also accumulate in FF, disturb COC physiology and compromise the oocyte’s maturation and developmental competence to generate an embryo [58]. For instance, NEFAs in concentrations associated with lipolytic conditions are classical examples of harmful molecules [59].

Given that the COCs are susceptible to concurrent pathological conditions (e.g. uterine infection, lipolytic disorders), it is important to unravel how specific molecules influence oocyte competence. Herein, we elaborated a relevant in vitro model to test the effects of a range of BHB concentrations on oocyte maturation and competence to develop to the blastocyst stage. With this goal, we first treated COCs with 0 (control), 2, 4 and 6 mM BHB during the entire period of in vitro maturation (∼22h) and evaluated the maturation rates based on the stereomicroscope visualization of an extruded first polar body. After observing over 500 oocytes (Control n:542; 2 mM BHB n: 529; 4 mM BHB n: 562; 6 mM BHB n: 525) from 11 biological replicates, we did not observe any detrimental effect (p=0.9331) of BHB at any dose on the oocyte’s capacity to undergo meiotic resumption during in vitro maturation (**Figure 6A**). Polar body extrusion rates were around ∼80% for all groups examined, indicating that, even in elevated levels (6 mM) of BHB, oocytes are able to resume and complete meiotic maturation. Moreover, as observed by stereomicroscopy, BHB-treated COCs possessed the same overall morphological appearance and cumulus expansion levels regardless of the dose utilized (**Figure 6B-G**). In contrast, exposure to NaBu, a closely related molecule and potent HDAC inhibitor, reduced maturation rates to around ∼35 % (p=0.0001) and affected the cumulus cell expansion (Supplemental Figure 8)

**Figure 6.**
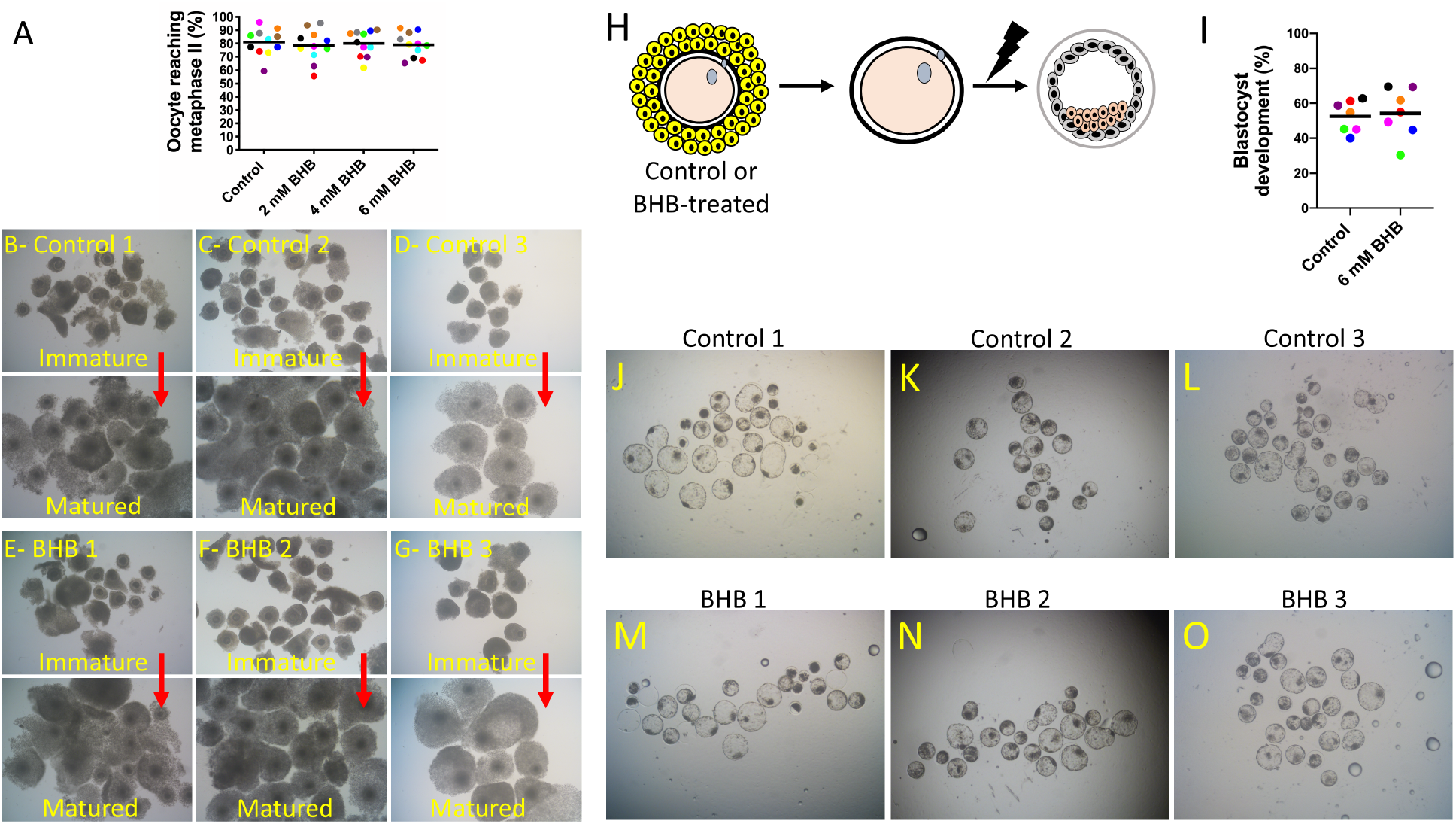
**(A)** Scatterplot showing the percentage of oocytes with extruded polar bodies (metaphase II arrest). Each color represents a biological replicate, p= 0.9331 from Tukey’s test. Brightfield pictures of COCs from control **(B-D)** or BHB-treated **(E-G)** group before and after (red arrow pointing) in vitro maturation. **(H)** Schematic illustration depicting the process of COC denudation, artificial activation of MII-arrested oocyte (lightning bolt), and in vitro culture for 7 days to generate the parthenotes. **(I)** The scatterplot shows the percentage of oocytes reaching the blastocyst stage (each color represents a biological replicate), p=0.5953 from Student’s t test. Brightfield pictures of Blastocyst-stage parthenotes (and few arrested-embryos) derived from the activation of oocytes matured previously under control conditions **(J-L)** or treated with 6 mM BHB **(M-O)**. All pictures were taken at 4x magnification and are from 3 different biological replicates.

To further evaluate the effects of BHB on oocyte competence, we artificially activated oocytes (**Figure 6H**) and cultured them for 7 days in vitro to produce parthenogenetic embryos (parthenotes). Since parthenogenesis involves the development of an embryo without paternal contribution, it allowed us to test whether BHB-exposed oocytes can reach the blastocyst stage relying only on their maternal-derived components [60]. Since no effects on in vitro maturation were observed at any BHB dose, we opted to evaluate the effect of BHB only at the highest concentration, i.e. 6 mM. Using 7 biological replicates, we activated 300 oocytes from the control group and 314 from the treated group. After 72 h of activation, cleavage rates did not differ between the control and 6 mM BHB (88.38±1.74 vs. 89.68± 3.99, p=0.5577). Moreover, after 7 days from activation, blastocyst development was similar between the control and treated groups (52.55 ± 3.43 vs. 54.30 ± 5.36; p=0.5953 (**Figure 6I-O**), indicating that the exposure of oocytes to BHB during in vitro maturation has no effect on the development of parthenotes to the blastocyst stage.

Our data showed that, even in elevated ketone concentrations found only in animals with severe clinical ketosis, the ability of oocytes to resume meiosis and develop to the blastocyst stage was not affected. It has been postulated that the ovulated oocyte has to pass by a whole period of follicular growth (∼90–100 days in the cow) under stress to ovulate with poor competence [61]. Thus, we cannot rule out the possibility that a more prolonged exposure or a different developmental window might compromise oocyte quality. Primordial, primary and secondary follicles seem to be more susceptible to stressors [61] than the pre-ovulatory follicles used here. However, previous studies have demonstrated that diets or pharmacological interventions as short as 4 days before ovulation can modify some oocyte features. The use of endoplasmic reticulum stress inhibitors reversed adverse effects of obesity and rescued the “metabolic health” on oocytes and restored embryo development, indicating that oocytes are sensitive to environmental cues even relatively close to the expected ovulation time [62].

Since the earliest stages of embryo development are primarily driven by oocyte components acquired during folliculogenesis, metabolic perturbations affecting the COC may affect preimplantational development. Indeed, maturation of oocytes for 24 h under high concentration of saturated NEFA has been shown to generate embryos with altered metabolism, i.e. reduced oxygen, pyruvate and lactate consumption [63]. Thus, preimplantation embryo metabolism can be preprogrammed during the final stage of oocyte maturation and affect some embryonic characteristics. Blastocysts can be classified based on their morphology, and their quality varies depending on criteria such as the number of cells, the trophectoderm to inner-cell mass ratio, the blastocoel expansion, the overall appearance, etc [64]. Herein we relayed only upon overall appearance to check the normalcy of the embryo and we did not observe noticeable alterations (**Figure 6J-O**). Albeit parthenogenetic embryos are inherently limited in their developmental capacity, we show no effects on developmental competence, indicating that the oocyte per si is not compromised after BHB exposure during in vitro maturation. In the future, it will be interesting to investigate whether the resulting embryos retain any metabolic, transcriptional, and epigenetic sequels caused by oocyte BHB exposure.

### BHB greatly increases H3K9bhb in cumulus cells but only faintly in oocytes during in vitro maturation

The oocyte growth phase itself is a protracted process that requires ∼3-4 months in cows and humans [61]. Histone PTMs change dynamically during the growth phase and play a crucial role in regulating transcription. At later stages, several histone PTMs are removed to ensure proper chromosome condensation and oocyte maturation while others are retained and play a role in early embryonic development [65]. During this growth period, oocytes and their surrounding cells are exposed for a very long time to environmental conditions (e.g. heat stress, metabolic disorders) that might affect their epigenetic marks [65]. Disturbed marks can lead to aberrant patterns of gene expression and compromise oocyte maturation and embryonic development, meaning that oocytes need to be shielded from stressors. Besides their nourishing role for the oocyte, cumulus cells also play a protective role to ensure that the oocyte is not affected by harmful compounds at advanced follicular stages [66]. The mechanisms of protection can be: (1) physical, by acting as a barrier between the oocyte and the extra follicular microenvironment, and (2) molecular, by metabolizing, degrading, accumulating or synthesizing enzymes or other biomolecules that in excess can jeopardize oocyte competence [67]. Recent reports indicate that COCs aspirated from cows in negative energy balance contain oocytes with altered patterns of DNA methylation when compared with heifers demonstrating that, in spite of being enclosed by cumulus, oocytes remain susceptible and undergo epigenome-wide alterations [68]. In vitro maturation of oocytes with NEFAs causes altered transcription in blastocysts, suggesting affected epigenetic marks [59].

Kbhb is a nutrient-sensitive epigenetic mark that has not been investigated in bovine oocytes so far. To seek for this mark, we exposed COCs to high concentration of BHB during the entire in vitro maturation (∼22h). First, we immunostained the whole COC (n>15) using an H3K9bhb antibody and quantified the immunoreactivity on nuclei (n>540) of cumulus cells (**Figure 7A-H**). As a result, intact COCs treated with 6 mM BHB increased the levels of Kbhb ∼3.61-fold in cumulus cells compared to control COCs (p<0.0001) (**Figure 7I**).

**Figure 7.**
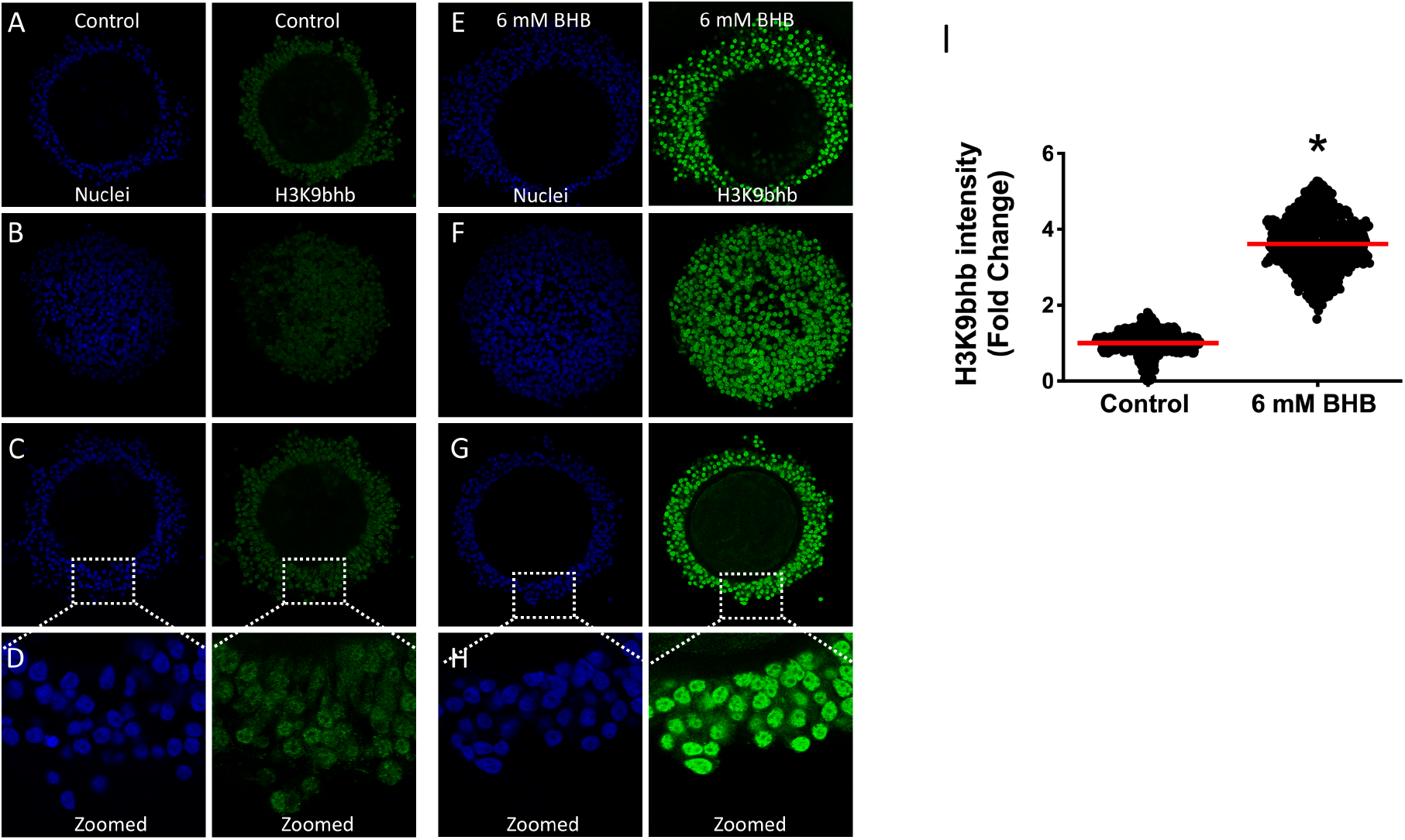
Immunostaining for H3K9bhb in 3 representative images of cumulus-oocyte complexes (COCs) from **(A-D)** Control and **(E-H)** 6 mM BHB treated groups. The images B (Control) and F (6 mM BHB) are a slice from the periphery of the COCs to illustrate that virtually all cells are enriched for Kbhb in treated oocytes. The images **D** and **H** are a cropped part from the images **C** and **G**, respectively, and they were manually zoomed only for illustrative purpose to show the details of cumulus cells staining. **(I)** Scatter plot showing H3K9bhb intensity in fold change (red dash represents the mean) in nuclei (n>540) of cumulus cells from COCs (n>15) measured by confocal microscopy. Asterisk (*) denotes statistical significance (p<0.0001) derived by Mann Whitney test. Nuclei were stained with Hoechst 33342 and H3K9bhb with Alexa Fluor 488 dye. Pictures were taken at 40x magnification.

Next, we removed the cumulus cells and focused our search for the effects of BHB on Kbhb levels in oocytes (**Figure 8A-H**). Curiously, after denuding the oocytes and staining for H3K9bhb we did not observe any Kbhb signal in oocyte metaphase plates (n=44) in the control group (**Figure 8A-D**). Regarding the treated group (n=43), the signal was only faintly detected in ∼53% of the oocytes analyzed (23/43) (**Figure 8E-H**). We repeated the experiment 4 times and also stained the oocytes in parallel with either H3K4me **(n=19,** **Figure 8I-L**) or H3K9ac **(n=14, Figure M-P)** to serve as a positive control for the immunostaining technique. As expected, all the oocytes showed immunoreactivity for H3K4me3 and H3K9ac. As a personal observation, the H3K4me3 possessed higher intensity compared to H3K9ac (i.e. we used very low laser power on the confocal microscope and observed strong signal), which is supported by previous studies that utilized immunostaining and quantified the overall abundance of these marks [65].

**Figure 8.**
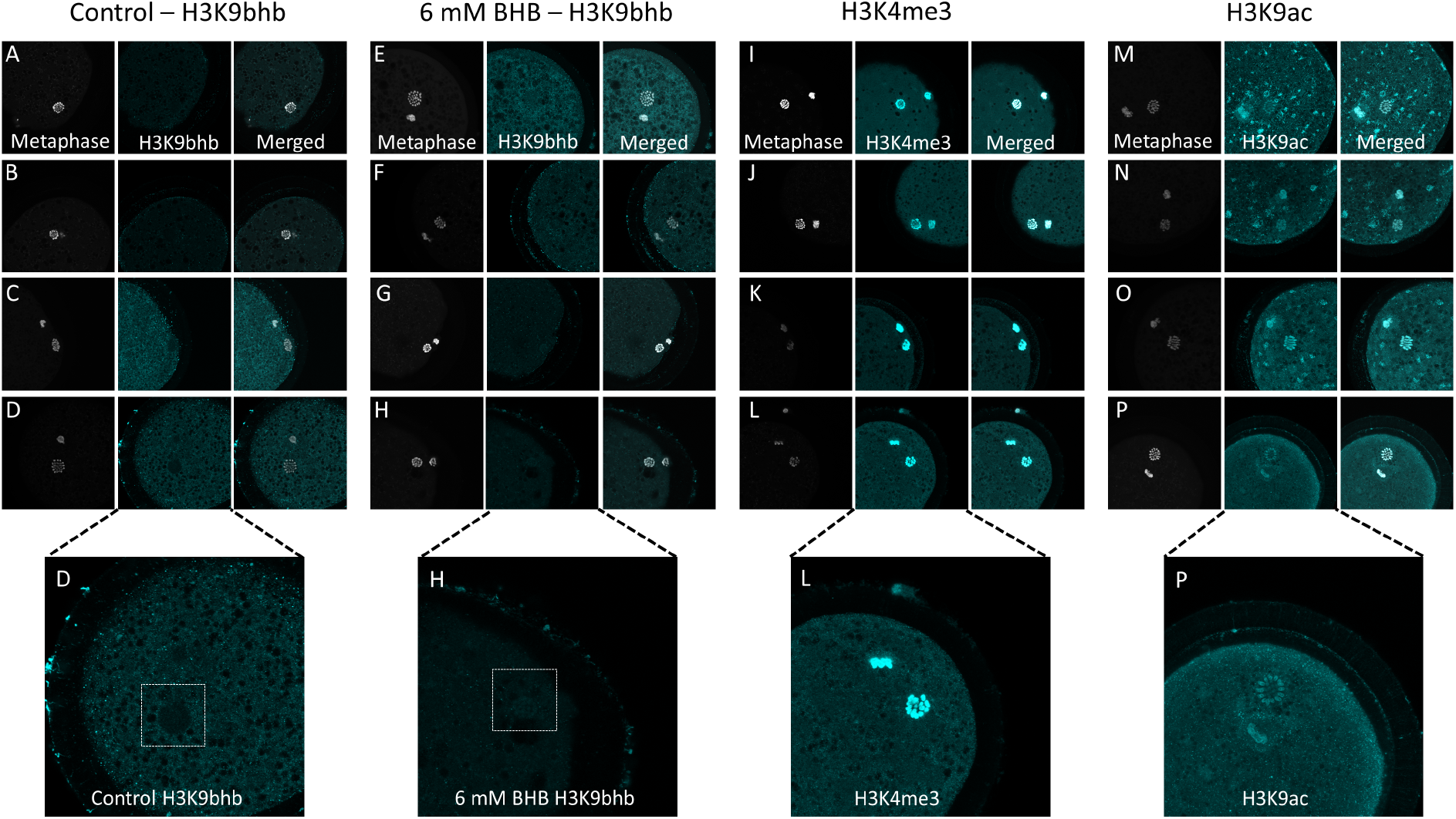
Representative pictures of 4 oocytes from control **(A-D)** and 6 mM BHB **(E-H)** groups stained for H3K9bhb. As positive controls, oocytes were stained also for H3K4me3 **(I-J)** and H3K9ac **(M-P)**. The pictures were taken in 100x magnification. The pictures **D**, **H**, **L**, and, **P** were enlarged for illustrative purpose, and the white dashed square used to highlight the metaphase plate area of Control and 6 mM BHB oocytes. Note that in the control group there is no evident signal, while in the BHB-treated it is possible to observe faint signal in the oocytes depicted in **G** and **H**. Cropped pictures highlighting only the metaphase plate can be found on Supplemental Figure 9. For control group were analyzed n=44 images; 6 mM BHB n=43; H3K4me3 n=19; H3K9ac n=14. Metaphase plates were stained with Hoechst 33342 and H3K9bhb with Alexa Fluor 488 dye. The metaphase plates were emulated for gray and the H3K9bhb staining for cyan to provide better contrast.

Several potential mechanisms could explain the close to absent levels of H3K9bhb signal on the metaphase plates of oocytes from COCs exposed to BHB during in vitro maturation. First, cumulus cells may be shielding the oocyte [69]. Supporting this view, it has been demonstrated that cumulus cells convert potentially toxic saturated into accumulated mono-unsaturated NEFAs [69, 70]. Later on, they store the fatty acids in lipid droplets, working as a “sponge”, and avoiding lipotoxicity due to excessive exposure of oocytes [71, 72]. Besides accumulation, another concomitant mechanism is the consumption of BHB as an energy source. Supplementation of granulosa cells with BHB increases cell numbers, suggesting its use as an energy and carbon source to proliferate [73]. Even if BHB is not completely consumed by cumulus cells, the mammalian oocytes possess around 200,000-400,000 copies of mtDNA per oocyte [74], which can contribute to BHB terminal oxidation and consequently less BHB available to be converted in its cognate CoA and tag histones. However, the synthetase responsible for BHB-CoA synthesis has not yet been discovered [34] and its presence in the oocyte’s cytoplasm remains elusive.

Second, BHB influx to the oocyte is controlled to avoid excessive levels. Albeit broadly accepted that BHB is a short fatty acid and can cross freely the membranes, its uptake seems to be controlled by the expression of transporters [19]. Supporting this view, the monocarboxylic acid transporters MCT1 and MCT2 carry BHB across the blood-brain barrier, and their expression can regulate brain BHB uptake [75]. Ketone body concentrations in human cerebrospinal fluid are considerably lower than that of blood during starvation [76]. The concentration of beta-hydroxybutyrate was 8.1 lower in the brain than in the arterial blood in adult and suckling rats [77]. Altogether, these data suggest that there are mechanisms regulating ketone uptake and permeability across tissues [23, 78]. It is not known whether limited tissue permeability to ketone bodies is peculiar to the brain or may be a general rule, for example limiting ketone permeability to mammalian eggs in stressful metabolic conditions. MCT1 mRNA is present throughout preimplantation development in mice [79], but it has not been demonstrated experimentally to be functional in oocytes. While the monocarboxylate transporter SLC16A6 is the key transporter for exporting BHB from the liver [80], the putative transporters that facilitate the uptake or its intracellular movement into target tissues remain to be identified. The use of isotope-labeled BHB may help solve the question of whether or to what extent circulating BHB reaches the oocyte’s cytoplasm.

Third, the erasers of Kbhb might be highly active in oocytes and hamper its accumulation. During oocyte growth and maturation, several histone PTMs change in abundance, reflecting their importance at given moments of this process. Some methylated and acetylated forms of histones (eg. H3K9ac, H4K12ac, H3K4me3 and H3K9me2) increase steadily throughout growth. However, during the final period of oocyte maturation (i.e. after germinal vesicle breakdown), as a general rule, the acetylated forms undergo widespread deacetylation [65], which happens to aid the process of chromatin compaction, a feature of the oocyte’s metaphase plate [48]. At this stage, the metaphase plate is characterized by chromatin highly compacted and with weakly detectable levels of marks associated with opening [65]. Because Kbhb deposition on histones affects charges and contributes to chromatin opening [81], it is highly likely they need to be removed at this period for chromatin closure. The widespread deacetylation wave is mainly caused by the HDAC2 activity [82]. Because HDAC1 and 2 were shown to be responsible for Kbhb removal [20], we speculate that Kbhb may be removed concomitantly with the deacetylation wave that occurs post germinal vesicle breakdown, leading to the faint or nearly absent signal observed in the oocytes. Other potential candidates to act as erasers that might contribute to Kbhb removal in oocytes are the sirtuins [3, 20]. Several members of the sirtuin family (e.g. Sirt1, 2, and 3) are present in oocytes where they potentially mitigate oxidative stress and delay oocyte aging [83]. Since BHB metabolism spare NAD+, which is a cofactor for sirtuins [6], excessive accumulation of Kbhb might be prevented due to enhanced activities of these enzymes caused by the availability of more co-factors. Supporting this idea, the SIRT1 and SIRT2 exhibited notable de-Kbhb activity towards core histones in vitro [20], and SIRT3 in HEK293 cells [44]. Moreover, we cannot exclude that changes in chromatin conformation may alter the accessibility of the epitope to the antibody and render them inaccessible to detect the mark. Also, based on mass spectrometry studies, Kbhb is far less abundant than Kac. The Kbhb showed low relative abundance ranging from 1 to –2%, as opposed to acetylation with global relative abundances between 15 and –30% in cells [33]. The low levels make its detection very difficult in cells undergoing massive deacylation such as maturing oocytes. Altogether, these concurrent hypotheses warrant further investigation to unravel the role of Kbhb on the oocyte’s epigenome.

### BHB treatment during IVM alters the transcriptome in cumulus cells

The transcriptome of granulosa and cumulus cells are known to be altered by several pathological (e.g. NEB in cows, Polycystic Ovarian Syndrome) and physiological conditions (e.g. aging, stages of follicular development) in mammals [84, 85]. Cumulus and mural granulosa cells reflect the characteristics of the oocyte and the follicular fluid, providing a noninvasive means to assess oocyte quality. Transcriptomic profiling of cumulus cells may help identify genes and pathways involved in oocyte and embryo competence [86]. Supporting this concept, cumulus cells from older women and livestock animals (i.e. mare, goat, cows) showed dysregulation of genes involved in cellular energy maintenance, antioxidant defense, mitochondrial function, apoptosis, and these alterations were associated with decreased embryo quality [87]. Granulosa cells collected from cows in severe negative energy balance showed downregulation of genes involved in cell proliferation, lipids and vitamin A/D metabolism [84]. Altogether, these data suggest that altered transcriptome may affect crucial functions in surrounding somatic follicular cells, such as by altering energy metabolism and communication, thereby affecting the meiotic resumption and oocyte competence [87]. We hypothesized that BHB alters the cumulus cell transcriptome and profiling them can aid to identify key genes and pathways involved in metabolic stress.

To this end, we carried out RNA-seq utilizing the Batch-Tag-seq, a robust method to profile gene expression data and investigate differentially expressed genes among samples. At least ∼11.79 millions of reads were obtained per sample, from which, on average ∼84.44% were uniquely mapped after sequencing. After mapping, we carried out Principal Component Analysis to further explore the data. As a result, we observed the formation of two clusters, containing 83% of the variance accumulated in the 2 principal components (**Figure 9A**), confirming that the control and BHB-treated samples are indeed distinct.

**Figure 9.**
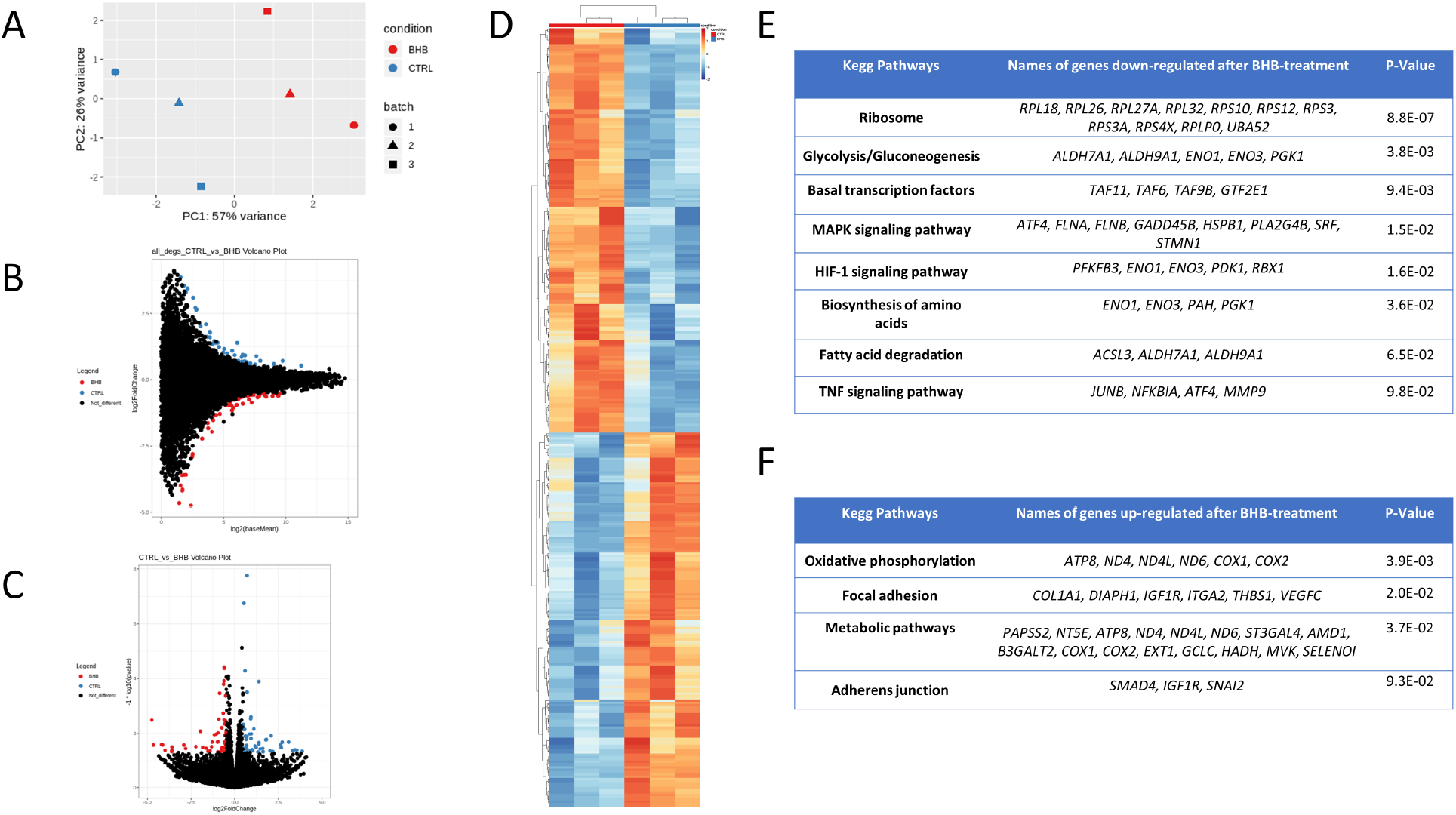
**(A)** Principal component analysis (PCA) graph showing the % of variance between samples from the Control versus BHB-treated group. **(B)** Smear Plot displaying the results of differentially expressed genes (DEG) after the RNA-seq analysis. **(C)** Volcano Plot displaying the results of differentially expressed genes (DEG) after the RNA-seq analysis. Red dots indicate the DEGs that were up-regulated in the BHB group, while blue dots represent the DEGs that were up-regulated in the control group. **(D)** Heatmap representing the variations in 345 differently expressed genes (DEGs) between cumulus cells from control group or treated with 6 mM BHB. Color scales vary from red (up-regulated genes) to blue (down-regulated genes). DEGs were considered to be either up-regulated or down-regulated when the adjusted p value <0.1. **(E)** Table showing the enriched Kegg Pathways and their respective genes obtained after submission of the list of genes downregulated after BHB treatment. **(F)** Table showing the enriched Kegg Pathways and their respective genes obtained after submission of the list of genes upregulated after BHB treatment.

Next, we performed differential expression analysis using DESEQ 2 where we identified 14459 genes (when considering at least 3 counts in at least 3 samples), from those 345 genes (2,38% of all detected genes) were differentially expressed between the groups based on an adjusted p value <0.1. From this total, 179 genes were down-regulated in BHB-group and 166 genes up-regulated compared to the control group. The whole gene expression profile can be observed in the smear and volcano plot below (**Figure 9B and 9C**). All differentially expressed genes are depicted in the heatmap (**Figure 9D**).

To gain insights about the biological pathways altered in the cumulus cells from COCs treated with BHB, we submitted the differentially expressed genes (DEGs) list to the David Bioinformatics Resource 6.8 platform [88] and carried out enrichment analysis. First, we submitted all the genes down-regulated in cumulus cells after treatment with BHB. We retrieved 8 Kegg pathways: 1-Ribosome; 2-Glycolysis/Gluconeogenesis; 3-Basal transcription factor; 4-MAPK signaling pathway; 5-HIF-1 signaling pathway; 6-Biosynthesis of amino acids; 7-Fatty acid oxidation (FAO); 8-TNF signaling pathway (**Figure 9 E**).

Based on the enriched pathways, our data suggests that BHB might be stimulating cumulus cells to adopt a parsimonious state and protecting against stressors by: (1) modulating several genes involved in Ribosome assembly and basal transcription factors that might reduce translation; (2) decreasing transcripts of a proliferative/stress-responsive (i.e. MAPK) and apoptotic pathway (i.e. TNF); (3) downregulating transcripts related mainly to glycolysis. BHB is interwoven with crucial mammalian metabolic pathways such as fatty acid oxidation (FAO), the tricarboxylic acid cycle (TCA), gluconeogenesis, de novo lipogenesis (DNL), and biosynthesis of sterols [19]. Moreover, BHB evolved to serve as an energy provider and energy saver molecule at the same time, and acts fine-tuning the control of lipid and glucose metabolism [16]. From an evolutionary point of view, these responses are aimed to help the organisms cope with nutrient deprivation. Decreasing glycolysis can aid to spare glucose for high-demanding organs such as the brain and the mammary gland from lactating cows.

While the treatment caused down-regulation of several genes responsible for glycolysis, submitting the list of genes up-regulated by BHB retrieved the following pathways: (1) Oxidative phosphorylation; (2) Focal adhesion; (3) Metabolic pathways; (4) Adherens junction (**Figure 9 F**).

During ketotic states BHB and the other ketone bodies become significant contributors to energy expenditure and are used in tissues rapidly. The fate of BHB in peripheral tissues is terminal oxidation in mitochondria to produce ATP. They are activated by mitochondrial thiolases into oxidizable form yielding two molecules of acetyl-CoA, which enter the TCA cycle [19]. Compared to fatty acid oxidation, BHB is more energetically efficient yielding more energy available for ATP synthesis per molecule of oxygen invested. Its oxidation keeps ubiquinone oxidized, raising the redox span in the electron transport chain and making more energy available to synthesize ATP [19, 89]. Additionally, BHB produces more heat during combustion compared to pyruvate, which increases the efficiency of ATP production from the mitochondrial proton gradient and reduces the production of free radicals. Curtailing the ROS production and thus oxidative stress might alter the activity of signaling networks that sense and respond to mitochondrial free radicals [89, 90].

Supporting all these data, the treatment of cumulus cells with BHB increased several genes related to oxidative phosphorylation, ketone utilization the mitochondrial respiratory complexes. Among the modulated genes, we observed an increase in HADH (Hydroxyacyl-Coenzyme A dehydrogenase), one of the thiolases involved in BHB activation to become readily oxidizable. Also, several members of NADH dehydrogenases (i.e. *ND4*, *ND4L* and *ND6*) were up-regulated. Indeed, ketone bodies have been demonstrated to enhance NADH oxidation in mitochondrial respiratory chain and consequently decreasing the reactive oxygen species in neurons [91]. Dysfunctions in some members of this family are implicated in metabolic disorders, including obesity, diabetes, and hypertension [92]. We observed that other members from the mitochondrial respiratory complexes were up-regulated by BHB treatment (i.e. *COX1*, *COX2* and *ATP8*). Alterations in the transcripts responsible for the metabolism can cause an increased in the mitochondrial efficiency and the net effect is a greater potential for ATP production.

Corroborating our findings, BHB or ketogenic diet have been shown to be effective in rescuing mitochondrial deficits in human diseases (e.g. Parkinson, MELAS) [93, 94]. Moreover, ketogenic diet produces low levels of redox signaling molecules, which in turn activate adaptive pathways and ultimately increasing the production of antioxidants such as glutathione (GSH) [95]. BHB increased the expression of *GCLC*, the catalytic subunit from the enzyme involved in the first and rate-limit step in GSH synthesis. It has been shown that glutathione (GSH) synthesized by intact cumulus cells during in vitro culture improved oocyte maturation and played an important role in fertilization and embryonic development [96]. Thus, BHB might be used as an additive in the maturation medium and its effect on GSH needs to be further investigated.

Besides the metabolic genes altered, we also observed a subset of genes involved in extracellular matrix (ECM) formation and angiogenesis. Curiously, several of these genes primarily cataloged as involved in angiogenesis processes (*VEGFC*, *SNAI2*, *THBS1*), are also involved in follicle development, oocyte competence and embryo development. A complete VEGF system (i.e. ligand and its receptors) was detected in bovine COCs. Exogenous VEGF supplementation during in vitro maturation improved embryonic development [97].

In the ovary, TGFβ-related proteins play crucial roles in controlling granulosa cell growth, differentiation, and steroidogenesis. SMAD4 is a central component of this pathway, and conditional knockout in COCs caused severe cumulus defects (e.g. abnormal expansion) and disrupted steroidogenesis [98]. Another crucial protein for granulosa cells’ function and consequently female fertility is IGF1R. Conditional knockdown of *IGF1R* causes sterility, ovaries smaller than control animals, no antral follicles even after gonadotropin stimulation and absence of oocytes in the oviduct after superovulation [99]. BHB treatment increased the expression of both these genes (*SMAD4* and *IGF1R*) corroborating our data of no noticeable effect on cumulus expansion and oocyte competence.

Another observation was that some genes involved in the extracellular matrix or cytoskeleton formation, adhesion, migration (i.e. *COL1A1*, *DIAPH1* and *ITGA2*) were up-regulated. The cumulus cell matrix is known to act: (1) as a scaffold to protect the oocyte while it is propelled into the oviduct, (2) to control the passage of small molecules including glucose and lipids, potentially altering the microenvironment that the oocyte is maturing, and (3) to attract, trap and interact with sperm at fertilization [49]. These mechanisms indicate that alterations in the matrix during maturation might last longer and affect embryo development. Another important observation is that, even if the cumulus cells are no longer communicating with the oocytes by TZPs at the final period of in vitro maturation, they still can send diffusible signals to the oocyte, oviduct and sperm [49]. Concluding, these genes altered in our RNA-seq identified specific genes and pathways altered by BHB on cumulus cell physiology and will provide entry points to carry out functional experiments aiming to improve fertility in cattle suffering metabolic disorders.

## CONCLUSIONS

In this article, we identified that histone lysine β-hydroxybutyrylation is present in bovine tissues in vivo and further confirmed that this epigenetic mark is responsive to BHB in vitro in a dose-dependent manner. We also demonstrated that COCs maturation with high concentrations of BHB did not affect the competence to complete meiotic maturation neither to develop to the blastocyst stage. BHB strongly induced H3K9bhb in cumulus, but this modification is only faintly detected in oocytes. In cumulus cells, BHB treatment mainly down-regulated genes involved in glycolysis and ribosome assembly and up-regulated genes involved in mitochondrial metabolism and oocyte development. The biology of histone β-hydroxybutyrylation remains largely unelucidated but is an area that should receive additional interest due to its potential relevance to gene expression reprogramming in response to metabolic stimuli [3–5,25,54,81]. A key goal for future work will be to understand mechanistically how BHB transmits signals from the environment to affect cellular functions and the bovine epigenome.

## MATERIALS AND METHODS

All chemicals and reagents used were purchased from Sigma-Aldrich Chemical Company (St. Louis, MO, USA) unless otherwise stated.

### Ethical Considerations

The present study was approved by the Animal Experimentation Ethics Committee of the University of São Paulo. The experiments were conducted in accordance with the International Guiding Principles for Biomedical Research Involving Animals (Society for the Study of Reproduction).

### Acid extraction of histones from cultured cells or tissues

Histones were extracted using the EpiQuick Total Histone Extraction Kit (Epigentek, Farmingdale, NY, USA, cat. # OP-0006) following the manufacturer’s instructions with few modifications. Dairy cow tissues were minced in tiny pieces utilizing a scalpel blade. Next, harvested cells or minced tissues were pelleted by centrifugation at 300 × g for 5 min at 4°C. Then, the cells were incubated in 200 μL of pre-lysis buffer for 10 min on ice, and centrifuged at 9,500 × g for 1 min at 4°C. The supernatant was removed, and the pellet was resuspended in 60 μL of lysis buffer and incubated for 30 min at 4°C. Samples were centrifuged at 13,500 × g for 5 min at 4°C. The supernatant, containing acid-soluble proteins, was transferred into a new vial and neutralized with 0.3 volumes of balance buffer supplemented with DTT. The proteins were quantified by Qubit 2.0 (Life Technologies) and stored in aliquots at -20°C.

### Coomassie staining

Two micrograms of acid-extracted histones were mixed with 4× Laemmli buffer (Bio Rad Laboratories, Hercules, CA, USA cat. #161-0747), denatured at 98°C for 5 min, and loaded into a sodium dodecyl sulphate–polyacrylamide gel. Proteins were fractionated by size on a 4%-15% SDS-PAGE run at a constant 100 V for 160 min. The gel was washed 3 times in ultra-pure water and stained with QC Colloidal Coomassie Stain (Bio Rad cat #1610803) for 20 h at RT. Next, the gel was washed for 3 h in ultrapure water to remove the background. The gel image was captured on a ChemiDoc MP Imaging System (Bio-Rad), and enrichment of histones was confirmed.

### Western Blotting for quantification of H3K9ac or H3K9bhb levels in somatic cells

To measure the relative Kac or Kbhb levels, 0.5 μg (to detect H3 or H3K9ac) or 3 μg of histones (H3K9bhb) were prepared and run as described above. Next, proteins were electroblotted onto nitrocellulose membranes (Bio-Rad) at a constant 80 V for 120 min using a wet transfer system (Bio Rad). The membranes were blocked with 3% BSA in Tris-buffered saline (TBS) + 0.1% of TWEEN (TBS-T) for 1 h at room temperature. The membranes were incubated overnight at 4°C under agitation with the primary antibody anti-H3K9ac (Sigma, cat # H9286) or anti-H3K9bhb (PTM Biolabs cat # PTM-1250) diluted 1:2,000 in blocking solution. Subsequently, membranes were washed 3 times with TBS-T for 5 min each and incubated for 1 h with peroxidase-conjugated anti-rabbit secondary antibody (Sigma, cat # A0545) diluted 1:10,000 in 1 % BSA solution in TBS-T. Finally, membranes were incubated for 30 seconds with Clarity Western ECL Substrate (Bio Rad cat. # 170-5060), and images were captured using the ChemiDoc MP Imaging System (Bio-Rad). Images were analyzed and bands were quantified using the software Image Lab 5.1 (Bio-Rad). The abundance of H3K9ac and H3K9bhb was calculated in relation to the loading control histone H3 antibody (Sigma, cat. # H0164; diluted 1:10,000) ran in parallel.

### Cell culture and treatment with Beta-hydroxybutyrate for confocal microscopy and western blot analysis

Bovine fibroblast cells were obtained as described previously [9]. Human Dermal Fibroblasts were purchased from Sigma (cat # 106-05A). A vial of cells was thawed, and plated at a density of 5 x 10^4^ cells per 35-mm Petri dish in cell culture medium (α-MEM (GIBCO BRL, Grand Island, NY, USA) supplemented with 10% (v/v) fetal calf serum (FCS) and 50 μg/ml gentamicin sulfate). At 24 h after plating, medium was replaced with fresh medium supplemented with:

-Bovine fibroblasts: Cultures were performed with 0 (control), 2, 4 and 6 mM BHB ((R)-(−)-3-Hydroxybutyric acid sodium salt, Sigma cat # 298360) or 5 mM Sodium Butyrate (NaBu, Sigma cat# 303410) for a period of 24 h.

-Human dermal fibroblasts: Cultures were performed with 0 (control), 2, 6 and 20 mM BHB for 24h.

At the end of cultures, cells were fixed with PFA 4% for immunostaining or tripsinized and recovered for histone extraction.

Regarding the experiment aimed at assessing Kbhb turnover, cells were either untreated (Control) or treated with 6 mM BHB (as described above), then washed 3 times with fresh culture medium and maintained without treatment for 0, 1, 2, 4 and 24 hours to monitor the Kbhb decay.

For the experiment involving inhibition of HDACs to increase Kbhb levels, cells were treated with 6 mM BHB, 5mM Nabu and 1 uM of TSA (Sigma cat # T8552) for 24 h.

### Immunostaining of fibroblasts, COCs and oocytes

Immunostaining was carried out as described previously [9], with few modifications. Samples were fixed in 4% paraformaldehyde (PFA) in PBS for 12 min, washed three times with PBS, and permeabilized with 1% Triton X-100 for 20 min. Samples were washed with PBS + 1 mg/ml BSA and blocked in PBS + 5% BSA + 0.3 M glycine solution for 1 h at RT. Next, samples were incubated overnight at 4°C on a rocking platform with the primary antibody anti-H3K9ac (1:1,000; Sigma, cat. # H9286) or anti-H3K9bhb (1:1,000; PTM biolabs cat. # 1250) or H3K4me3 (1:1,000; Abcam cat. # ab8580) diluted in PBS + 3% BSA + 0.1% Tween 20 solution. After extensive washing (5 x for 10 min) with PBS + 0.1% Tween 20, samples were incubated with the secondary antibody Alexa Fluor 488-conjugated goat anti-rabbit IgG (Life Tech, cat. # A11008) or Alexa Fluor 647-conjugated goat anti-rabbit IgG (Life Tech, cat. # A21244) diluted in PBS + 3% BSA + 0.1% Tween-20 at 1:1,500 for 1 h at room temperature protected from light under agitation. Appropriate negative controls were obtained by substituting the primary antibody for a rabbit polyclonal IgG isotype control (Abcam, Cambridge, MA, USA, cat # AB27478) or omitting the primary antibody (secondary-only control). Samples were counterstained with Hoechst 33342 (1μg/ml) and then mounted on a glass slide in Prolong Gold Antifade Mountant (Life Tech, cat. # P36935).

Samples were visualized and images were captured using a confocal microscope (TCS-SP8 AOBS; Leica, Soims, Germany), and the same settings were used for all images within the same experiment. All samples for a given experiment were processed for immunostaining together, and all images were taken at the same laser power, thereby enabling direct comparison of signal intensities. The pixel intensities of the images were analyzed using the software ImageJ (National Institutes of Health, Bethesda, MD, USA). Nuclei were individually outlined using the manual tool, and the average signal intensity was calculated.

### Oocyte recovery and in vitro maturation

Ovaries were collected from a local slaughterhouse and transported to the laboratory in an insulated container filled with saline solution (0.9% NaCl) at ∼30°C. Ovaries were washed several times and kept in pre-warmed saline solution during the ovary aspiration. Oocytes were aspirated from visible antral follicles (∼2 to 8 mm) using an 18 G needle connected to a syringe. Cumulus-oocyte complexes (COCs) containing compact cumulus cell layers and homogeneous cytoplasm were selected and matured in groups of 50 COCs in 500 μl of M199 with Earle’s salts, L-glutamine, 2.2 g/L sodium bicarbonate (GIBCO BRL, Grand Island, NY, USA cat. # 11150-059), 10 µg/mL FSH (Vetoquinol), 2 µg/mL β-estradiol (Sigma cat # E2758), 50 μg/mL gentamicin sulfate, 0.2 mM sodium pyruvate, and 10% FBS. In vitro maturation (IVM) were performed for ∼22 h in a humidified atmosphere of 5% CO_2_ in air at 38.5°C.

### Production of parthenogenetic embryos

After IVM, matured oocytes had their cumulus cells removed by incubation in 400 µL of trypsin (Tryple Express) and gentle pipetting. The denuded oocytes were washed with M199 containing 25 mM HEPES (Gibco, cat. # 12350-039) and artificially activated by incubation with 5 μM ionomycin diluted in M199 supplemented with 1 mg/mL BSA. Next, presumptive zygotes were washed 3 times in M199 + 30 mg/mL of BSA and incubated in SOF medium containing 2 mM 6-DMAP for 3 h. Then, presumptive zygotes were extensively washed and cultured in SOFaa [100] for 7 d in a humidified atmosphere of 5% CO2 in air at 38.5 °C.

### Treatment of cumulus-oocyte complexes (COCs) with BHB during in vitro maturation

In experiments involving COC treatment, Control group was matured using regular maturation medium as described above. As for the treated group, a solution of BHB was added to the maturation media freshly prepared. The experiment where the COCs were treated during the firsts 8 h, IVM medium was initially supplemented with 2 mM BHB and oocytes kept for 4 h. After this, an additional volume of BHB (+2 mM) was added, totalizing 4 mM and the oocytes were kept for another 4 h. Next, cumulus cells were removed by pipetting and histones acid extracted to measure H3K9bhb.

To evaluate maturation rates, COCs were treated with 0, 2, 4 and 6 mM BHB or 5 mM NaBu for 22 h, denuded and examined for the presence of the first polar body using stereomicroscopy.

As for experiments involving RNA-seq and staining of intact COCs and oocytes, COCs were treated for 22 h with 6 mM BHB or not (Control). At the end of treatment, COCs were: (1) fixed for staining and measurement of H3K9bhb, or (2) denuded and the cumulus cells utilized for RNA-seq and, (3) oocytes were stained to investigate the presence of H3K9bhb on metaphase plate.

### RNA extraction and library preparation

Three pools of cumulus cells from COCs matured or not in the presence of 6 mM β-hydroxybutyrate containing ∼150,000 cells were used for RNA-seq. Total RNA was isolated using the Qiagen RNeasy Mini Kit, checked for purity using automated electrophoresis (all samples presented RQI >9.8), and then submitted to the Genome Center at the University of California, Davis for library preparation and sequencing according to vigorously standardized protocol in the facility.

Gene expression profiling was carried out using a 3’ Tag-RNA-Seq protocol. Barcoded sequencing libraries were prepared using the QuantSeq FWD kit (Lexogen, Vienna, Austria) for multiplexed sequencing according to the recommendations of the manufacturer (Lexogen). The fragment size distribution of the libraries was verified via micro-capillary gel electrophoresis on a Bioanalyzer 2100 (Agilent, Santa Clara, CA). The libraries were quantified by fluorometry on a Qubit fluorometer (LifeTechnologies, Carlsbad, CA), and pooled in equimolar ratios. Up to forty-eight libraries per lane were sequenced on a HiSeq 4000 sequencer (Illumina, San Diego, CA). The sequencing was carried by the DNA Technologies and Expression Analysis Core at the UC Davis Genome Center, supported by NIH Shared Instrumentation Grant 1S10OD010786-01.

### RNA-seq pipeline

The data were processed in the Laboratory of Molecular Genetics and Bioinformatics (LGMB) of the Medical School of Ribeirão Preto, University of São Paulo (FMRP-USP). All data passed through a quality check by FASTQC software (http://www.bioinformatics.babraham.ac.uk/projects/fastqc). Reads were filtered according to data quality with the software TRIMMOMATIC [101], with 40 phred quality, minimum, at start/end of the read, medium of 40 phred quality and minimum read size of 50 bps. After quality filtering we repeated QC analysis with FASTQC software. Samples that passed quality screening were then aligned with the *Bos taurus* genome (assembly ARS-UCD1.2), using Rsubread [102] with standard parameters. Alignment quality was then verified with a final report generated using the MULTIQC software [103]. After alignment the reads were counted using the feature counts [104].

### Differential gene expression analysis

First we performed a principal component analysis (PCA), followed by the differential gene expression analysis using DESEQ2 [105] R package considering an adjusted p values <0.1. For gene expression values representation, we used the variance normalization transformation (vsd, “varianceStabilizingTransformation” function from DESEQ2 package). For enrichment analysis we submitted the lists of differentially expressed genes to David Bioinformatics Resources 6.8 [88].

### Statistical analysis

Statistical analysis was performed using the GraphPad Prism 5 (GraphPad Software, San Diego, California, USA). Data were tested for normality of residuals and homogeneity of variances using the Shapiro-Wilk test and analyzed as follows. Data from experiments involving: western blot and immunostaining in bovine and human fibroblasts were analyzed by ANOVA followed by Tukey’s post hoc test or Kruskal-Wallis’ test. Data from experiment involving western blot in cumulus cells was analyzed by Student’s test. Data from experiments using immunostaining in COCs was analyzed by Mann Whitney test. Frequency data (experiments involving oocyte maturation and parthenogenetic embryonic developmental rates) was analyzed using the chi-squared test. Differences with probabilities of p < 0.05 were considered significant. In the text, values are presented as the means or the means ± the standard error of the mean (s.e.m.). All experiments were repeated at least three times unless otherwise stated.

## Supplementary data

Supplementary data are available online.

## Data availability

Data are available upon request.

## Author Contributions

**J.R.S.**, **P.J.R.**, and **F.V.M.** designed the research; **J.R.S.**, **R.P.N.**, **M.C.**, and **R.V.S.** performed the experiments; **J.R.S.**, and **R.P.N.** analyzed the results; **J.R.S.**, **F.P.**, **J.C.S.**, **P.J.R.**, **L.C.S.**, and **F.V.M.** interpreted the results; **J.R.S.**, **L.C.S.**, and **F.V.M.** wrote the paper.

## Acknowledgments

Authors thank the slaughterhouse for ovaries’ donation. Members from Laboratory of Molecular Morphophisyology and Development for technical assistance and critical reading.

---

**Supplemental Figure 1.**
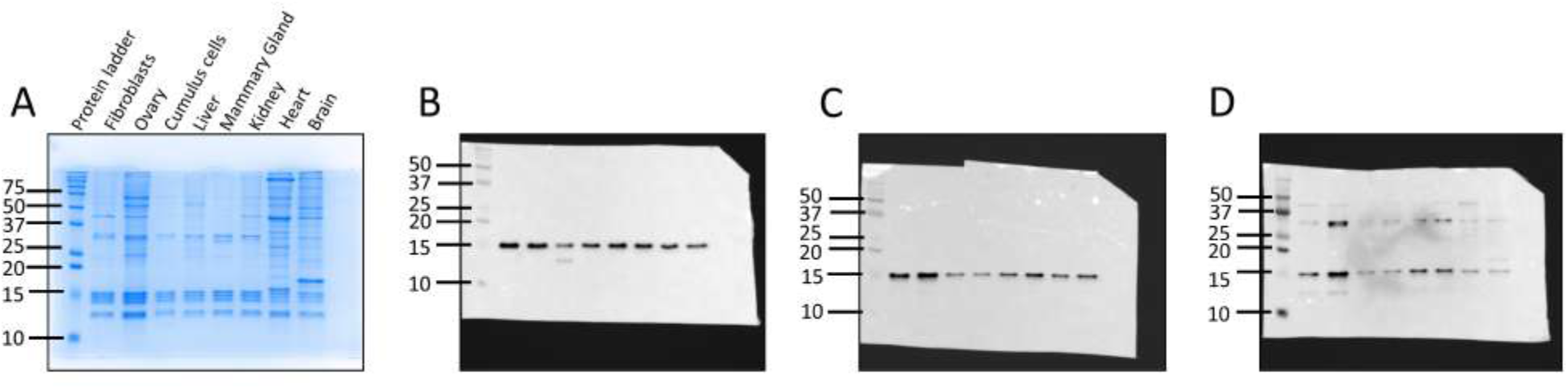
The full gel/blots stained with coomassie blue **(A)** and immunoblots for total H3 **(B)**, H3K9ac **(C)** and H3K9bhb **(D)** from dairy cow cells and tissues. The immunoblots were merged with the colorimetric picture showing the protein standards used to estimate the molecular weight of proteins. Numbers on the left represent the molecular weight in kDa.

**Supplemental Figure 2.**
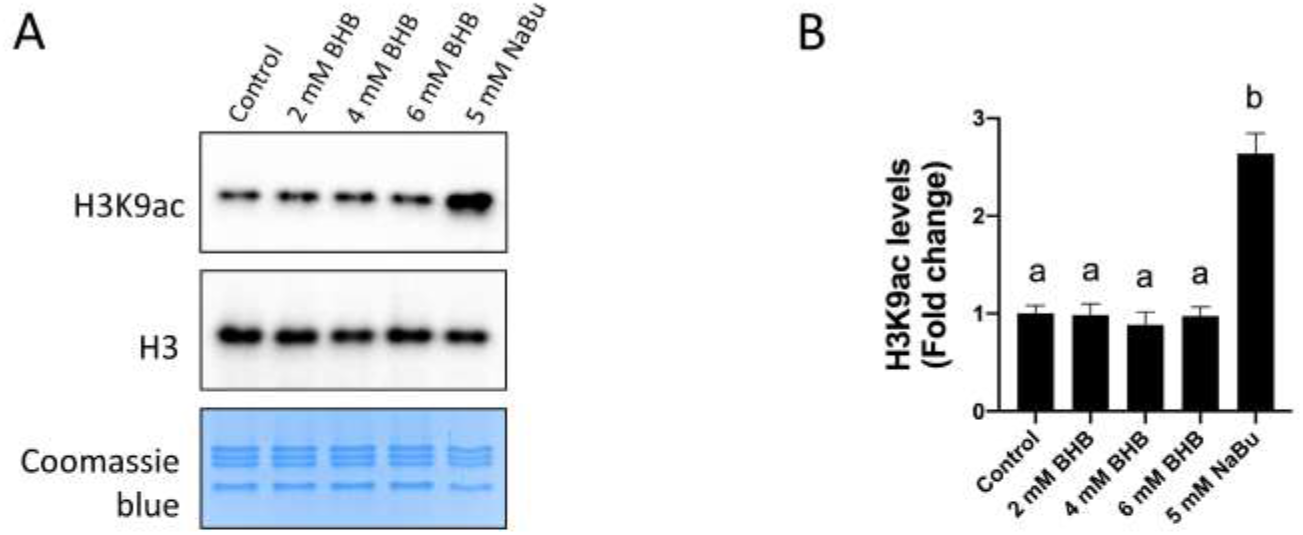
**(A)** Representative gel stained with coomassie blue and immunoblots for H3K9ac and total H3 from bovine fibroblasts cultured as described in the manuscript. **(B)** Bar graph shows the H3K9ac levels in fold change (± sem) measured by Western blot (WB). Treated group values are represented as fold change compared to the control group. Different letters indicate statistical differences between groups as measured by Tukey’s test (p<0.0001).

**Supplemental Figure 3.**
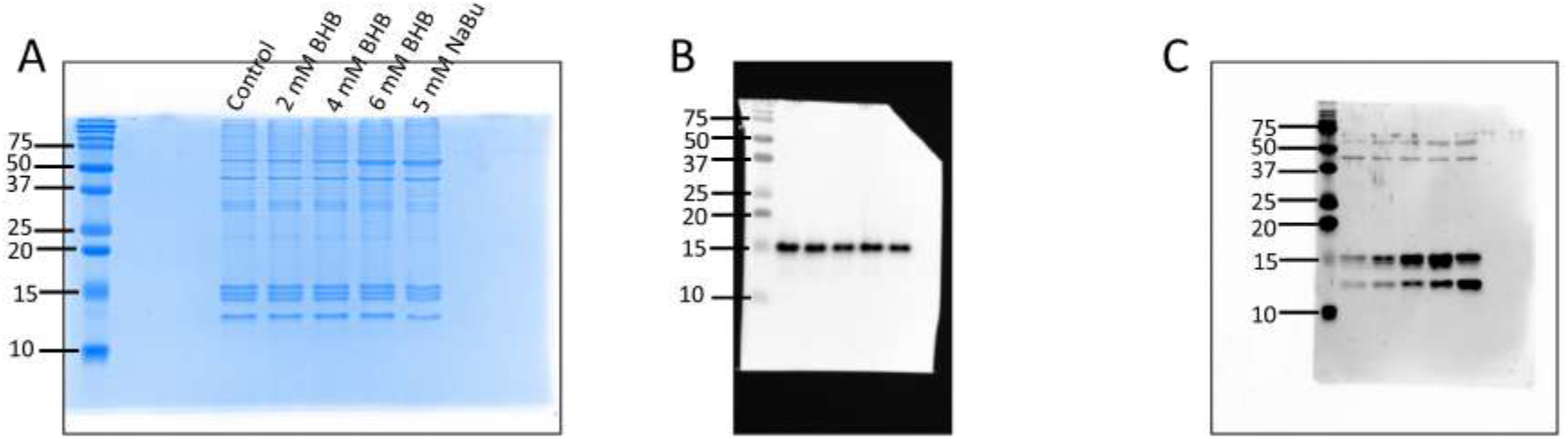
The full gel/blots stained with coomassie blue **(A)** and immunoblots for total H3 **(B)** and H3K9bhb **(C)** from bovine fibroblasts. The immunoblots were merged with the colorimetric picture showing the protein standards used to estimate the molecular weight of proteins. Numbers on the left represent the molecular weight in kDa.

**Supplemental Figure 4.**
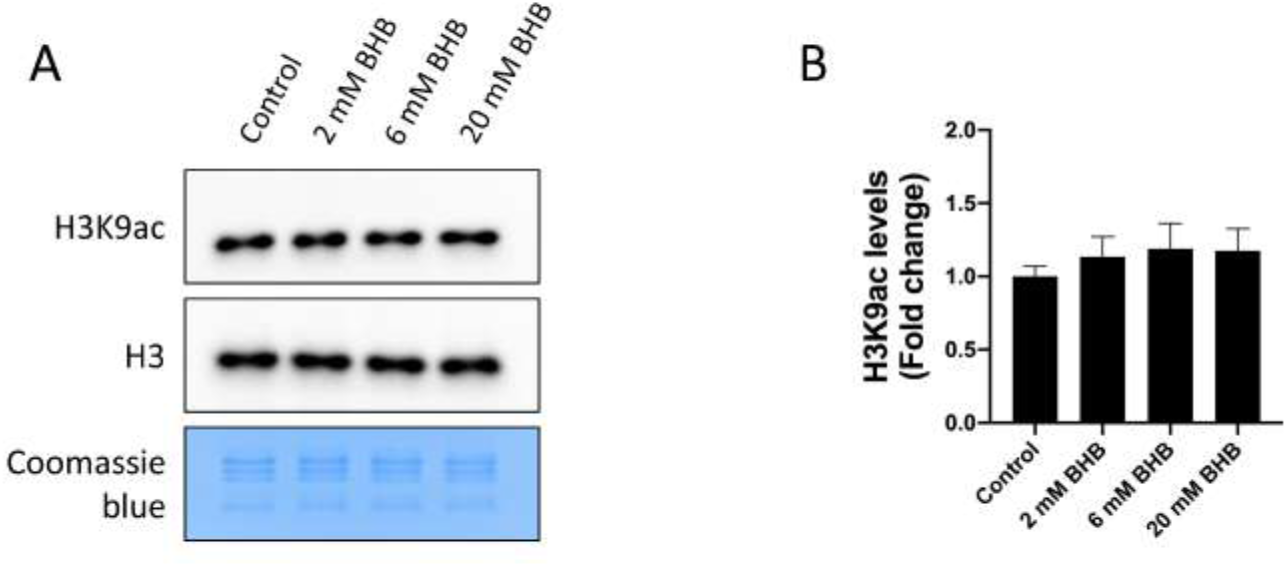
**(A)** Representative gel stained with coomassie blue and immunoblots for H3K9ac and total H3 from human dermal fibroblasts cultured as described in the manuscript. **(B)** Bar graph shows the H3K9ac levels in fold change (± sem) measured by Western blot (WB). Treated group values are represented as fold change compared to the control group. No statistical differences between groups as measured by Tukey’s test (p=0.7684).

**Supplemental Figure 5.**
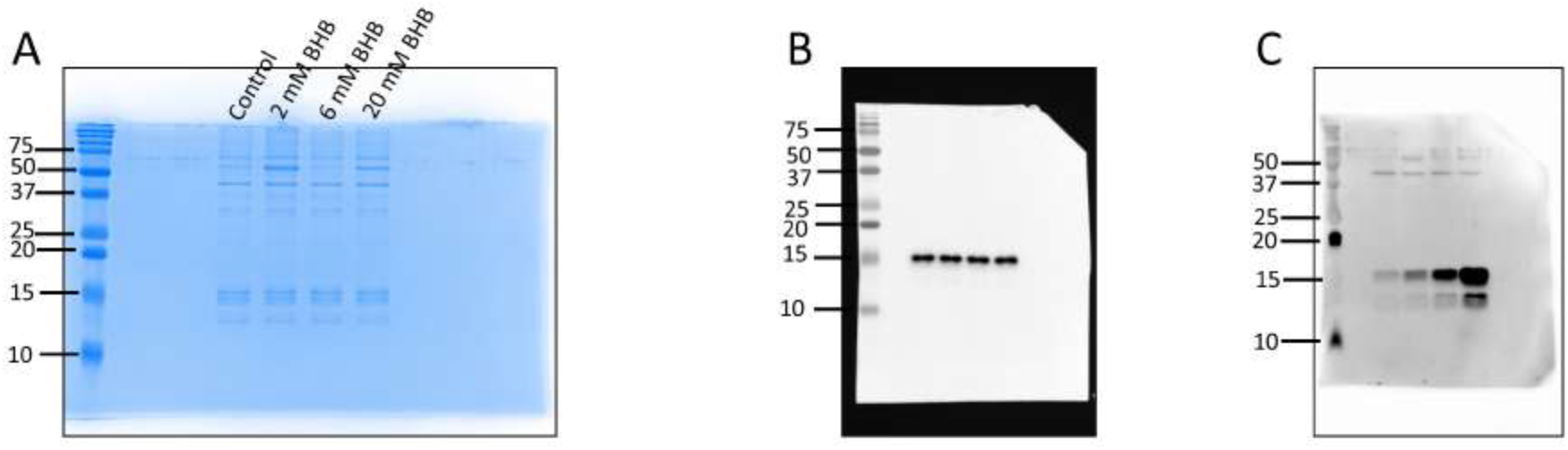
The full gel/blots stained with coomassie blue **(A)** and immunoblots for total H3 **(B)** and H3K9bhb **(C)** from human dermal fibroblasts. The immunoblots were merged with the colorimetric picture showing the protein standards used to estimate the molecular weight of proteins. Numbers on the left represent the molecular weight in kDa.

**Supplemental Figure 6.**
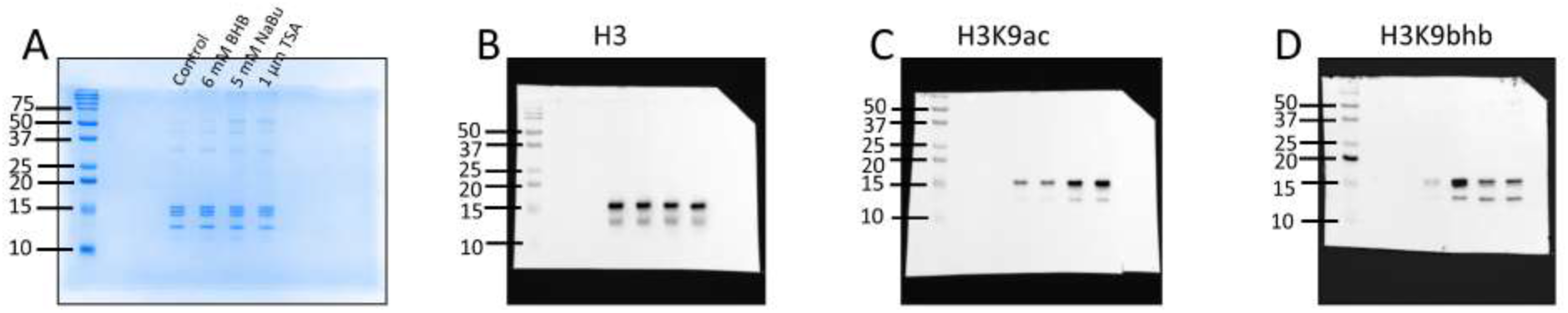
The full gel/blots stained with coomassie blue **(A)** and immunoblots for total H3 **(B)**, H3K9ac **(C)** and H3K9bhb **(D)** of bovine fibroblasts from control and treated groups exposed to 6 mM BHB, 5 mM NaBu and 1 µM TSA. The immunoblots were merged with the colorimetric picture showing the protein standards used to estimate the molecular weight of proteins. Numbers on the left represent the molecular weight in kDa.

**Supplemental Figure 7.**
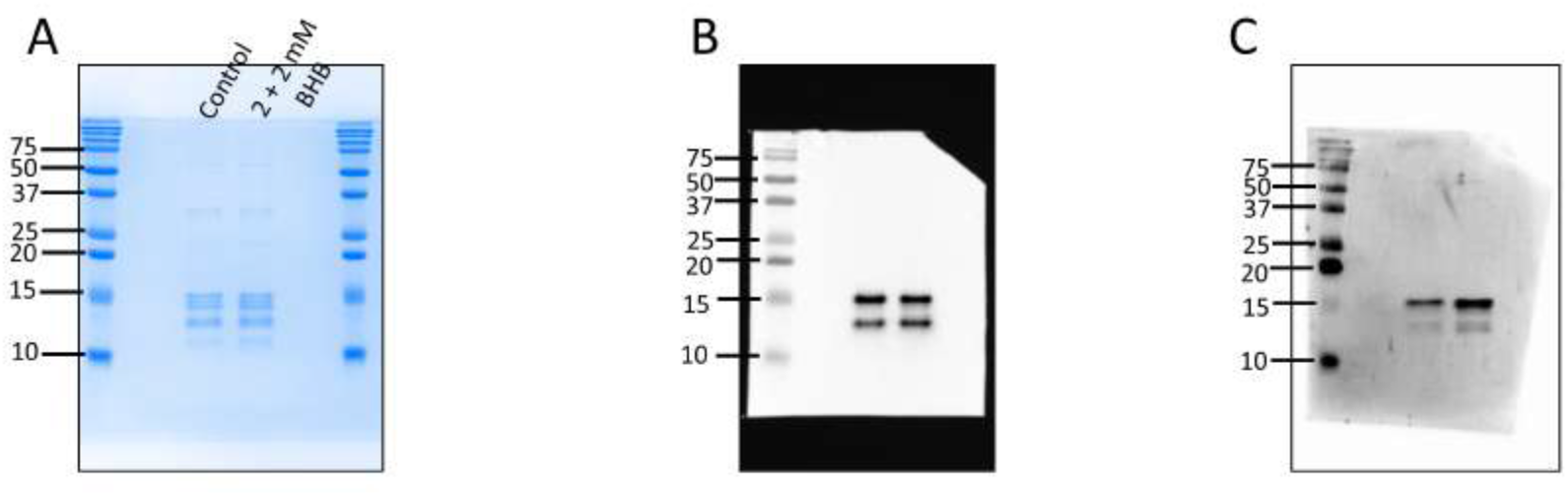
The full gel/blots stained with coomassie blue **(A)** and immunoblots for total H3 **(B)** and H3K9bhb **(C)** from cumulus cells cultured for 8 h in IVM medium supplemented with 2 to 4 mM BHB or not (Control). The immunoblots were merged with the colorimetric picture showing the protein standards used to estimate the molecular weight of proteins. Numbers on the left represent the molecular weight in kDa.

**Supplemental Figure 8.**
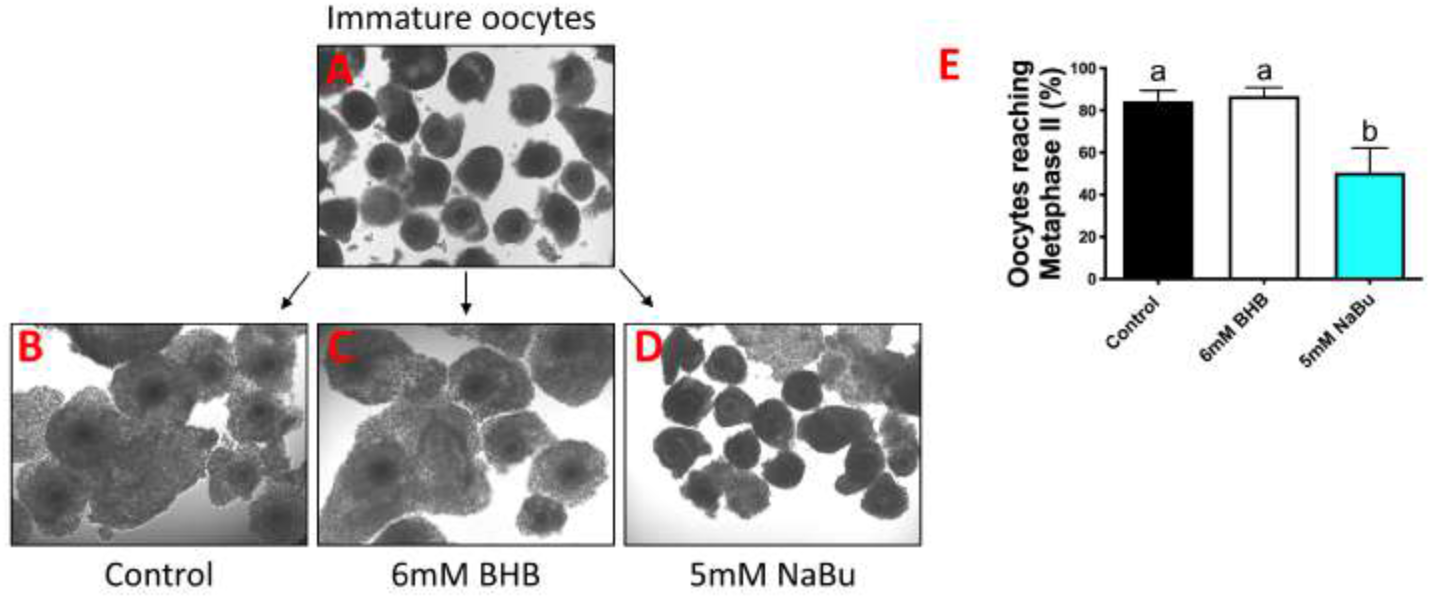
Brightfield pictures of immature COCs **(A)** and after maturation from control **(B)**, 6 mM BHB **(C)**, and 5 mM NaBu **(D)** groups. Note that COCs matured with NaBu failed in properly expand the cumulus cells. **(E)** The bar graph shows the percentage of oocytes with extruded polar bodies (metaphase II arrest). Different letters indicate statistical differences (p=0.0001) using Tukey’s test. All pictures were taken at 4x magnification.

**Supplemental Figure 9.**
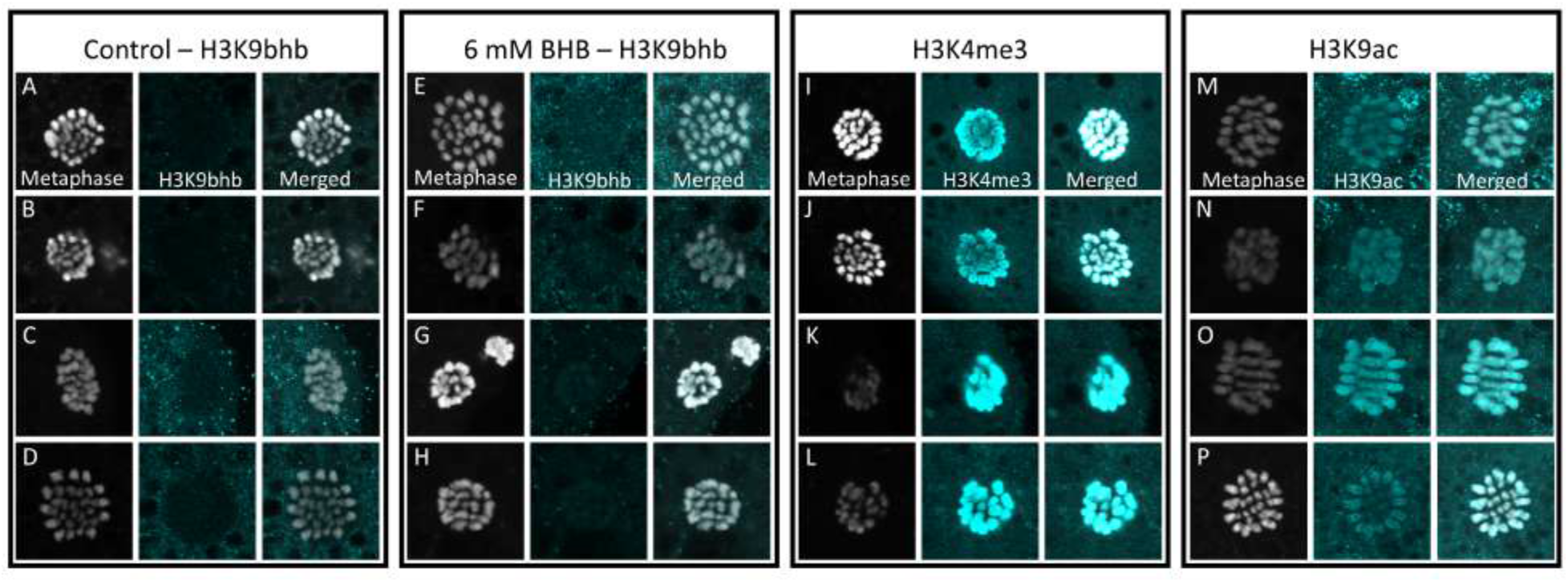
Cropped pictures of oocyte metaphase plates depicted in the **Figure 8** from control **(A-D)** and 6 mM BHB **(E-H)** groups stained for H3K9bhb. As positive controls, oocytes were stained also for H3K4me3 **(I-J)** and H3K9ac **(M-P)**. The pictures were taken in 100x magnification. The images were cropped for illustrative purpose and to highlight only the metaphase plate area. Metaphase plates were stained with Hoechst 33342 and H3K9bhb with Alexa Fluor 488 dye. The metaphase plates were emulated for gray and the H3K9bhb staining for cyan to provide better contrast.

